# DNA-dependent phase separation by human SSB2 (NABP1/OBFC2A) protein points to adaptations to eukaryotic genome repair processes

**DOI:** 10.1101/2023.09.15.557979

**Authors:** Zoltán J. Kovács, Gábor M. Harami, János Pálinkás, Natalie Kuljanishvili, József Hegedüs, Hajnalka Harami-Papp, Lamiya Mahmudova, Lana Khamisi, Gergely Szakács, Mihály Kovács

**Affiliations:** ELTE-MTA “Momentum” Motor Enzymology Research Group, Department of Biochemistry, Eötvös Loránd University, Pázmány P. s. 1/c, H-1117 Budapest, Hungary; HUN-REN–ELTE Motor Pharmacology Research Group, Department of Biochemistry, Eötvös Loránd University, Pázmány P. s. 1/c, H-1117 Budapest, Hungary; HUN-REN Institute of Molecular Life Sciences, Research Centre for Natural Sciences, Hungarian Academy of Sciences, Magyar Tudósok körútja 2, H-1117 Budapest, Hungary; Center for Cancer Research, Medical University of Vienna, Borschkegasse 8a, 1090 Wien, Austria

## Abstract

Single-stranded DNA binding proteins (SSBs) are ubiquitous across all domains of life and play essential roles *via* stabilizing and protecting single-stranded (ss) DNA as well as organizing multiprotein complexes during DNA replication, recombination, and repair. Two mammalian SSB paralogs (hSSB1 and hSSB2 in humans) were recently identified and shown to be involved in various genome maintenance processes. Following our recent discovery of the liquid-liquid phase separation (LLPS) propensity of *E. coli* (Ec) SSB, here we show that hSSB2 also forms LLPS condensates under physiologically relevant ionic conditions. Similar to that seen for EcSSB, we demonstrate the essential contribution of hSSB2’s C-terminal intrinsically disordered region (IDR) to condensate formation, and the selective enrichment of various genome metabolic proteins in hSSB2 condensates. However, in contrast to EcSSB-driven LLPS that is inhibited by ssDNA binding, hSSB2 phase separation requires single-stranded nucleic acid binding, and is especially facilitated by ssDNA. Our results reveal an evolutionarily conserved role for SSB-mediated LLPS in the spatiotemporal organization of genome maintenance complexes. At the same time, differential LLPS features of EcSSB and hSSB2 point to functional adaptations to prokaryotic *versus* eukaryotic genome metabolic contexts.

## INTRODUCTION

Single-stranded DNA-binding proteins (SSBs) are crucial for genome maintenance and DNA replication in all organisms. They recognize and bind single-stranded (ss) nucleic acids, stabilizing and protecting them from harmful chemical alterations and nucleolytic cleavage. Besides replication protein A (RPA), the major nuclear ssDNA-binding protein, two new mammalian SSB proteins, SSB1 (NABP2/OBFC2B) and SSB2 (NABP1/OBFC2A), have recently been discovered (1,2). Both paralogs have been shown to be involved in DNA repair, B cell and embryonic development (1,3–9). Besides their roles in genome maintenance, their capability to bind ssRNA also raises their possible involvement in RNA metabolic processes (6,10–12). Unlike human (h) SSB1, which is broadly expressed, hSSB2 expression is tissue-specific, similar to what was also observed for mouse orthologs (2). According to the Human Protein Atlas database (http://www.proteinatlas.org/), moderate hSSB2 mRNA levels were detected in lymphoid tissues, spleen, the gastrointestinal tract, as well as the male and female reproductive system, and low expression levels are indicated elsewhere.

Available data indicate the functional involvement of hSSB2 in DNA repair and the response to various forms of genomic stress (5,6,9,13,14). A mouse model with a deleted mouse (m) SSB2 gene is viable with no obvious phenotype under stress-free conditions (15). In contrast, the absence of mSSB1 results in perinatal death and developmental abnormalities (5). However, a partial functional overlap between the two proteins is evident, as simultaneous deletion of both genes is embryonic lethal and loss of either SSB1 or SSB2 upregulates the expression of the other paralog (6,9). Importantly, SSB2 appears to promote cell viability and DNA repair in some contexts. It colocalizes with pH2AX upon DNA damage (16), and its ablation leads to rapid cell death in wild-type mouse embryonic fibroblasts (5) and increased sensitivity to DNA damaging agents in HeLa cells (13). Upon human T-lymphotropic virus type 1 (HTLV-1) infection, suppressed hSSB2 expression is linked to defects in cell proliferation (17). Furthermore, a role for SSB2 in the protection of newly synthesized telomeres was also proposed (5,16,18).

In line with their multifaceted roles, hSSB1 and hSSB2 have been shown to interact with a wide array of proteins involved in both nuclear and cytoplasmic functions (1,13,19–22). Either hSSB1 or hSSB2 can interact with the INTS3 and INIP proteins to form the SOSS complex, which is involved in the recognition and repair of double-stranded (ds) DNA breaks (12,13,23–25). However, hSSB1 appears to have a more pronounced role in the initiation of homologous recombination-based DNA double-strand break repair, whereas SSB2 exerts a less prominent role in ATM kinase activation (13) and checkpoint arrest (23,26) during the response to gamma irradiation-elicited DNA damage. Recent data also indicate distinct functional specialization for hSSB paralogs. hSSB2 is involved in the nucleotide excision repair pathway (NER) and can recognize and rapidly localize to cyclobutene pyrimidine dimers formed upon UV irradiation (14,27), whereas hSSB1 is selectively involved in the base excision repair pathway due to its unique ability to recognize oxidized guanine bases (28–30).

Tight nucleic acid binding and the capability for diverse protein-protein interactions are mediated by the N-terminal oligonucleotide-oligosaccharide binding (OB) domain and the C-terminal intrinsically disordered region (IDR) of SSB proteins, respectively (**Fig. 1A-B**) (10,31). The OB fold is highly conserved, with only a few amino acid substitutions between SSB1 and SSB2 (1) and, accordingly, both proteins share a common mechanism for ssDNA binding (27). Recently, we showed that the IDR present in *E. coli* (Ec) SSB has an additional function as it mediates homotypic SSB condensation leading to liquid-liquid phase separation (LLPS) (32). The similar architecture of bacterial and human SSB homologs raises the possibility that hSSB2 is also capable to form protein condensates via its IDR, in line with our previous *in silico* predictions (32). A number of eukaryotic, prokaryotic, and viral proteins have been shown to form protein condensates, often together with nucleic acids, mediated by multivalent intermolecular IDR-IDR interactions (33–36). In the cell, LLPS appears to play a critical role in regulating the spatiotemporal distribution and activity of protein complexes— among other functions, allowing for rapid, adaptive response to various forms of stress (33–36). Accordingly, a role for EcSSB condensation was implicated in genomic stress response (32). EcSSB localizes to the bacterial cell membrane in multiple foci, consistent with the *in vitro* LLPS propensity of the protein in the absence of nucleic acids (37). However, ssDNA binding dissolves EcSSB condensates (32) and, under genomic stress leading to ssDNA exposure, the protein quickly relocates from the membrane to affected genomic loci (37). Human RPA also undergoes LLPS mediated by an N-terminal IDR present in the RPA-B subunit (38). Interestingly, RPA LLPS is triggered by ssDNA binding, and *in vivo* RPA condensation is associated with telomere clustering (38).

**Figure 1.**
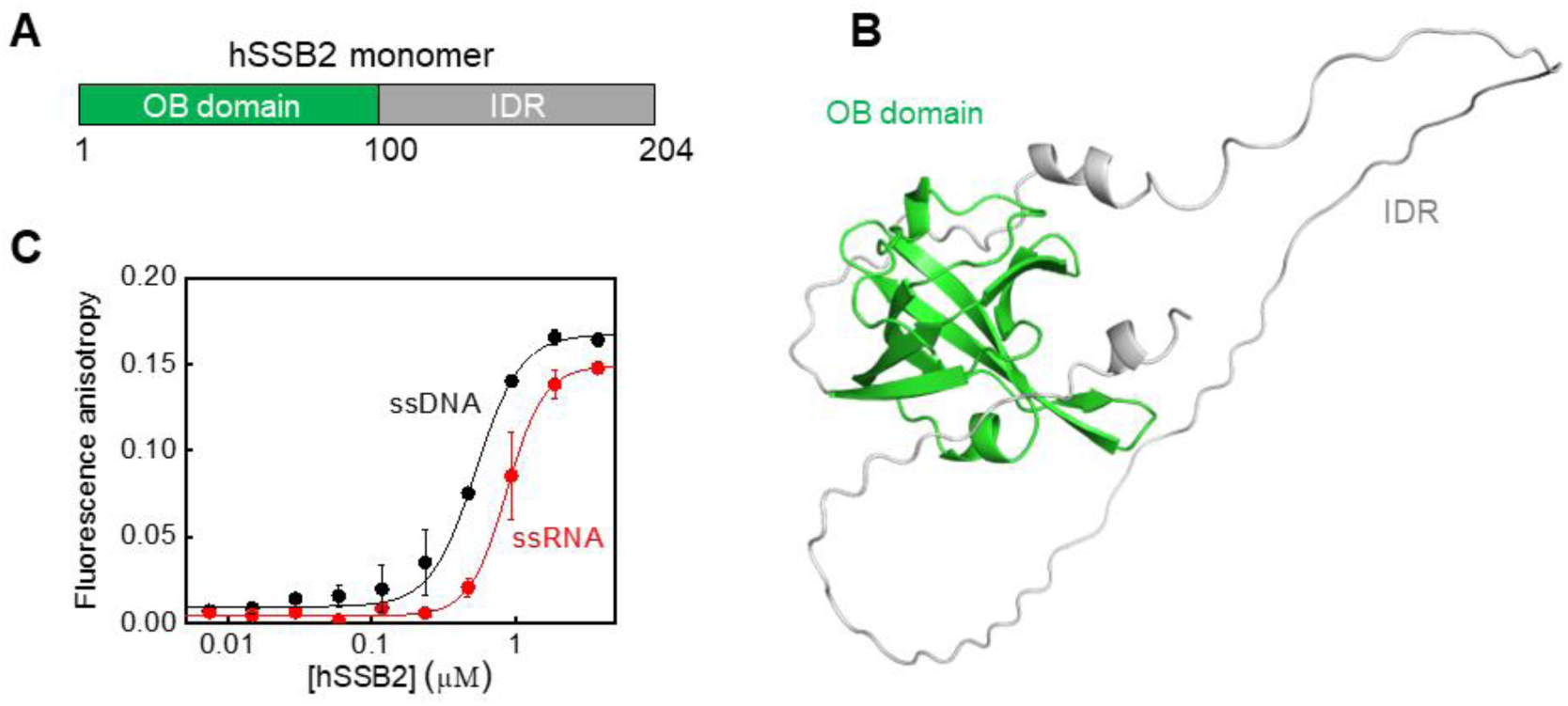
Human single-stranded DNA-binding protein 2 (hSSB2), comprising an oligosaccharide-oligonucleotide binding (OB) domain and an intrinsically disordered region (IDR), binds ssDNA and ssRNA with similar affinities. **(A)** Schematic of the domain structure of hSSB2. Numbers indicate amino acid positions at the boundaries of the oligosaccharide-oligonucleotide binding (OB) domain and the intrinsically disordered region (IDR). **(B)** Predicted 3D structure of hSSB2 (AlphaFold Protein Structure Database ID: Q96AH0). **(C)** Fluorescence anisotropy measurements performed by titrating 10 nM 3’-fluorescein labeled, 36-mer ssDNA or ssRNA with increasing concentrations of hSSB2. Solid lines show best fits using the Hill equation. Best-fit parameter values are reported is **Supplementary Table 1**. Means ± SEM are shown for *n* = 3.

Extending the scope of the above observations, here we show that hSSB2 can form phase-separated, liquid-like coacervates with ssDNA *in vitro*, and that hSSB2’s protein interaction partners are selectively enriched in the condensates. The process is reversible and inhibited by 1,6-hexanediol and supraphysiological concentrations of chloride ions, indicating a *bona fide* LLPS process. Condensation characteristics are also influenced by redox environment, as LLPS particles appear under reducing conditions, whereas oxidation appears to solidify hSSB2-ssDNA coacervates. hSSB2 phase separation is mediated by intermolecular IDR-IDR interactions and our data indicate a scaffolding function for ssDNA that provides multivalency of protein-protein interactions. Interestingly, ssRNA has a markedly lower LLPS-inducing effect on hSSB2 than ssDNA, indicating nucleic acid type-specific effects in addition to the scaffolding function. Taken together, our results demonstrate the capability of hSSB2 for condensate formation in a cellular context upon ssDNA binding. Moreover, in our recent work we have shown that hSSB1 also undergoes nucleic acid-dependent phase separation, albeit with properties that differ from those of hSSB2 described herein (see Discussion) (39). Thus, phase separation emerges as common feature among IDR-containing SSB proteins, with specific properties shaped by varying evolutionary demands in bacterial and eukaryotic nucleic acid metabolic contexts.

## RESULTS

### hSSB2 binds ssDNA and ssRNA and undergoes liquid-liquid phase separation upon ssDNA binding

Our previous *in silico* analysis indicated a high LLPS propensity for both hSSB2 and hSSB1 conferred by their C-terminal IDR region (32). Thus, we aimed to test our prediction experimentally using purified recombinant hSSB2. C-terminally His-tagged hSSB2 (referred to as hSSB2 in this article) was purified from *E. coli* cells using the method of Vidhyasagar et al. (10). Surprisingly, during our initial purification attempts, highly concentrated hSSB2 fractions (>300 µM; SSB concentrations are expressed in monomer units throughout the manuscript) became opaque and showed high turbidity, indicating either protein aggregation or LLPS. Importantly, the process was reversible, as the protein solution turned transparent upon addition of 100 mM MgCl_2_. Based on this effect, hSSB2 aliquots (**Fig. S1**) were subsequently stored in 100 mM MgCl_2_ to preserve the soluble form.

Next, we measured hSSB2’s ssDNA and ssRNA binding affinity *via* a fluorescence anisotropy-based method (**Fig. 1C**), using fluorescein-labeled 36-mer ssDNA and ssRNA oligonucleotides. In line with previous findings (10), hSSB2 binds both types of ss nucleic acid, evidenced by the protein concentration-dependent increase in the fluorescence anisotropy of the oligonucleotides. These results were also corroborated by electrophoretic mobility shift assays (**Fig. S2A**). Analysis of binding isotherms (**Figs. 1C** and **S2B**) using the Hill equation revealed a slight preference for ssDNA over ssRNA and a cooperative binding behavior (**Table S1**). In addition, the observed cooperativity and the formation of distinct hSSB2-nucleic acid complexes in EMSA experiments suggest simultaneous and cooperative interaction of multiple hSSB2 molecules with the applied 41-nt oligonucleotides and/or a protein concentration and DNA binding-dependent change in the oligomerization state of hSSB2. We note that fluorescence anisotropy assays indicated a higher affinity and lower cooperativity value for both nucleic acid types compared to the EMSA results (**Table S1**). This variation may stem from technical differences, as the EMSA is not a genuine equilibrium method, unlike the fluorescence anisotropy assay. Complex dissociation during electrophoresis may lead to underestimation of binding density (40). In addition, if multiple complex forms with varying dissociation rates are present, the same effect could result in a higher apparent cooperativity, as weaker complexes become less represented. Nevertheless, the above experiments confirm hSSB2’s functionality in binding either ssDNA or RNA, indicate a moderate preference for ssDNA over ssRNA, and suggest hSSB2 oligomerization upon nucleic acid binding, in line with previous results (10).

To assess whether hSSB2 is capable of phase separation with or without ssDNA, we tested the presence of visible particles in SSB solutions containing moderate salt concentrations (25 mM Tris-HCl pH 7.4, 50 mM KCl, 10 mM MgCl_2_, 1 mM DTT; this condition, termed LLPS buffer, is used as standard condition unless otherwise stated) using differential interference contrast (DIC) microscopy (**Fig. 2A**). In the absence of ssDNA, the hSSB2 solution was transparent and only a small number of amorphous particles were observed, likely originating from impurities and/or limited protein aggregation. Turbidity experiments also indicated the presence of a limited number of particles scattering light (**Fig. S3**). However, upon addition of ssDNA (dT79, 79-mer homo-deoxythymidine), the solution turned markedly more turbid, and we observed the appearance of numerous spherical particles (**Fig. 2A**). Fluorescence microscopy experiments using a mixture of unlabeled and fluorescently labeled hSSB2 (see Methods) confirmed that the observed droplets are formed by the protein upon addition of ssDNA (**Fig. 2A**). ssDNA concentration-dependent fluorescence microscopy experiments using either fluorescently labeled hSSB2, or labeled ssDNA together with unlabeled hSSB2, corroborated that the hSSB2 protein forms coacervates with ssDNA (**Fig. 2B**). Condensate formation was enhanced even by low, substoichiometric amounts of ssDNA (**Fig. 2B**). The extent of condensation increased with increasing ssDNA concentration (**Fig. 2C-D**) until reaching saturation at a stoichiometry of about 2.5-5 mol hSSB2 protein per mol dT79 molecule. Higher ssDNA concentrations suppressed the extent of condensation, but droplets were still observed at a 4-times excess of ssDNA over hSSB2 (**Fig. 2B-D**).

**Figure 2.**
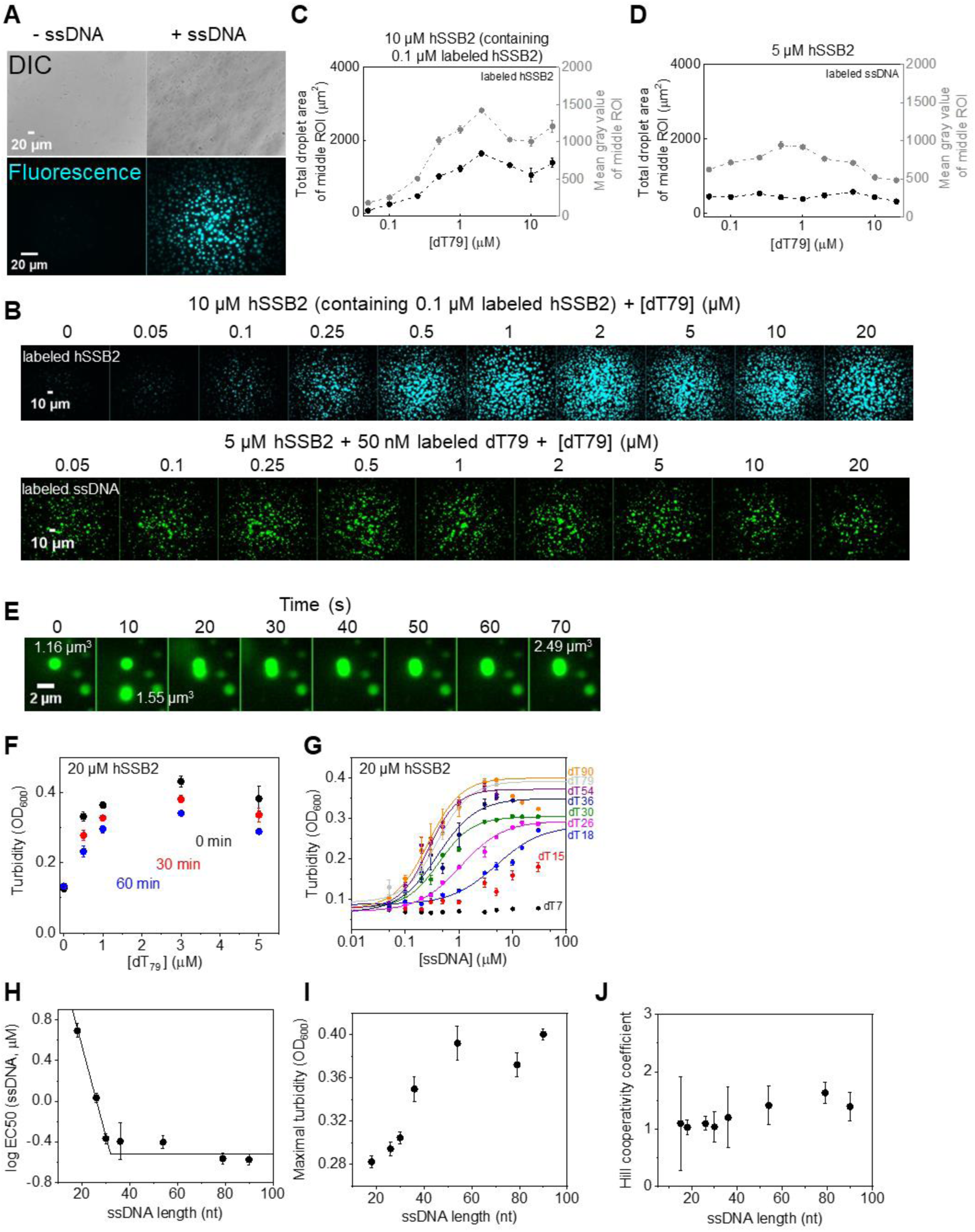
hSSB2 forms phase-separated condensates in the presence of ssDNA. **(A)** Top row: DIC images of 10 µM unlabeled hSSB2 in the presence and absence of 1 µM ssDNA (79-mer homopolymer deoxythymidine, dT79). Bottom row: fluorescence microscopic images of 10 μM hSSB2 (containing 0.1 µM fluorescein-labeled hSSB2) in the presence and absence of 1 µM dT79. Samples were incubated for 60 minutes before imaging. **(B)** Fluorescence microscopic images showing the ssDNA concentration dependence of hSSB2 droplet formation in LLPS buffer. Upper row: images obtained upon 60-min incubation of 10 μM hSSB2 (containing 0.1 μM fluorescein-labeled hSSB2) and indicated concentrations of dT79. Bottom row: images obtained upon 30-min incubation of 5 μM hSSB2, 50 nM Cy3- labeled dT79 and indicated concentrations of dT79 **(C-D)** dT79 concentration dependence of the total droplet area and mean gray values determined from the middle section (see Methods) of fluorescence images shown in panel **B**. Means ± SEM are shown for *n* = 3. **(E)** Spherical droplet morphology and volume additive fusion indicate liquid nature of hSSB2 condensates. 5 µM hSSB2 was mixed with 2 µM dT79 containing 0.1 µM Cy3-dT79, and fluorescence images were obtained at 10-s intervals. Indicated droplet volumes were calculated based on the diameter of droplets. **(F)** Post-mixing time and dT79 concentration dependence of the turbidity of solutions containing 20 µM hSSB2 and indicated concentrations of dT79. Means ± SEM are shown for *n* = 3. **G)** ssDNA (oligo-dT) length and concentration dependence of the turbidity of solutions containing 20 µM hSSB2 and the indicated concentrations of oligo-dT molecules of varying length. Means ± SEM for *n* = 6 (*n* = 3 for dT7) are shown. Solid lines show best fits using the Hill equation. Best-fit values are shown in panels **H-J**. **(H-J)** ssDNA length dependence of (**H**) the logarithm of ssDNA concentration required for half saturation (log EC50 (ssDNA)), (**I**) maximal turbidity values and (**J**) Hill coefficients obtained in experiments shown in panel **G**. Error bars show fitting standard errors. Solid line in panel **H** shows best fit using **Equation 1** (see Methods). Best-fit parameters ± fitting errors obtained were: length of saturating ssDNA (*b*) = 32.0 ± 0.9 nt, y intercept (c) = 2.31 ± 0.23 and amplitude (A) = −2.84 ± 0.24.

The liquid nature of hSSB2-ssDNA coacervates was also corroborated by frequently observed rapid droplet fusion events (**Fig. 2E**). Moreover, 1,6-hexanediol, a widely used substance capable of dissolving LLPS condensates (41), inhibited droplet formation in a concentration dependent manner (**Fig. S4**). Turbidity measurements further confirmed that ssDNA binding enhances phase separation and indicated rapid condensation kinetics as the turbidity of hSSB2 solutions reached its maximal level concomitantly with manual mixing with ssDNA (**Fig. 2F**). Condensates remained stable over prolonged incubation times, but the turbidity signal decreased slightly with time (**Fig. 2F**), likely due to fusion of droplets (**c.f. Fig. 2E**), similar to that proposed for *Ec*SSB (32).

We tested the ssDNA length dependence of hSSB2 condensation by titrating hSSB2 (20 µM) with increasing concentrations of oligo-dT molecules of various length in turbidity experiments (**Fig. 2G**). The shortest ssDNA molecule tested, dT7, was unable to trigger LLPS even up to 30 µM concentration. However, dT15 and longer ssDNA molecules increased the turbidity of the hSSB2 solution in a length and concentration dependent manner (**Fig. 2G**). In case of dT79 and dT90, high ssDNA concentrations decreased the turbidity, in line with microscopy results (**Fig. 2C-D**). In case of shorter ssDNA strands, such a decrease was not detected in the applied concentration regime. The ssDNA concentration required for half-saturation (as revealed by fits using the Hill equation) decreased with increasing ssDNA length, and remained constant above 32 ± 1 nt (**Fig. 2H**). Maximal turbidity values also increased with ssDNA length, but showed saturation at greater lengths (**Fig. 2I**). At greater ssDNA lengths, mild cooperativity (Hill coefficient ∼1.4) in condensate formation was observed (**Fig. 2J**). Taken together, the above results show that ssDNA binding to hSSB2 induces the formation of hSSB2-ssDNA coacervates, and efficient condensation requires a minimum ssDNA length of 32 nucleotides.

### Condensation is more pronounced with ssDNA than ssRNA, and is mediated by the IDR region

The relatively high ssRNA binding affinity of hSSB2 (**Figs. 1C and S2, Table S1**) raised the question whether hSSB2 condensation is triggered by ss nucleic acids, regardless of their type. To test this possibility, first we monitored hSSB2 droplet formation in fluorescence microscopy experiments using increasing concentrations of either dT41 ssDNA or U41 ssRNA, containing a fixed amount of their respective fluorescently labeled variants (**Fig. 3**). Fluorescent particles were observed both with ssDNA and ssRNA even at the lowest tested nucleic acid concentration (**Fig. 3A**). However, it must be noted that these particles may originate from binding of the labeled strands to aggregates/preassembled condensates existing already in the SSB2 samples prior to the addition of nucleic acids (cf. **Fig. 1A** and **Fig. S3**). Importantly, the extent of condensation increased with dT41 ssDNA concentration and, after reaching a maximum, it started to decrease (**Fig. 3B-C**) similarly to that observed with dT79 (**Fig. 2B-D**). Contrary, ssRNA did not increase the extent of condensation markedly (**Fig. 3B-C**), and at high ssRNA concentrations particles completely disappeared (**Fig. 3A**), either due to disassembly of condensates upon ssRNA binding or due to the competition between unlabeled and labeled ssRNA for hSSB2. Although the application of labeled hSSB2 protein instead of labeled oligonucleotides resulted in discernible ssRNA-induced phase separation, ssDNA-induced LLPS was more robust even under these conditions, corroborating ssDNA specificity of hSSB2 LLPS (**Fig. 3A, D-E**). Results obtained at lower salt conditions excluded the possibility that the lack of pronounced LLPS stimulation by ssRNA could originate from altered sensitivity to ionic conditions (**Fig. S5**).

**Figure 3.**
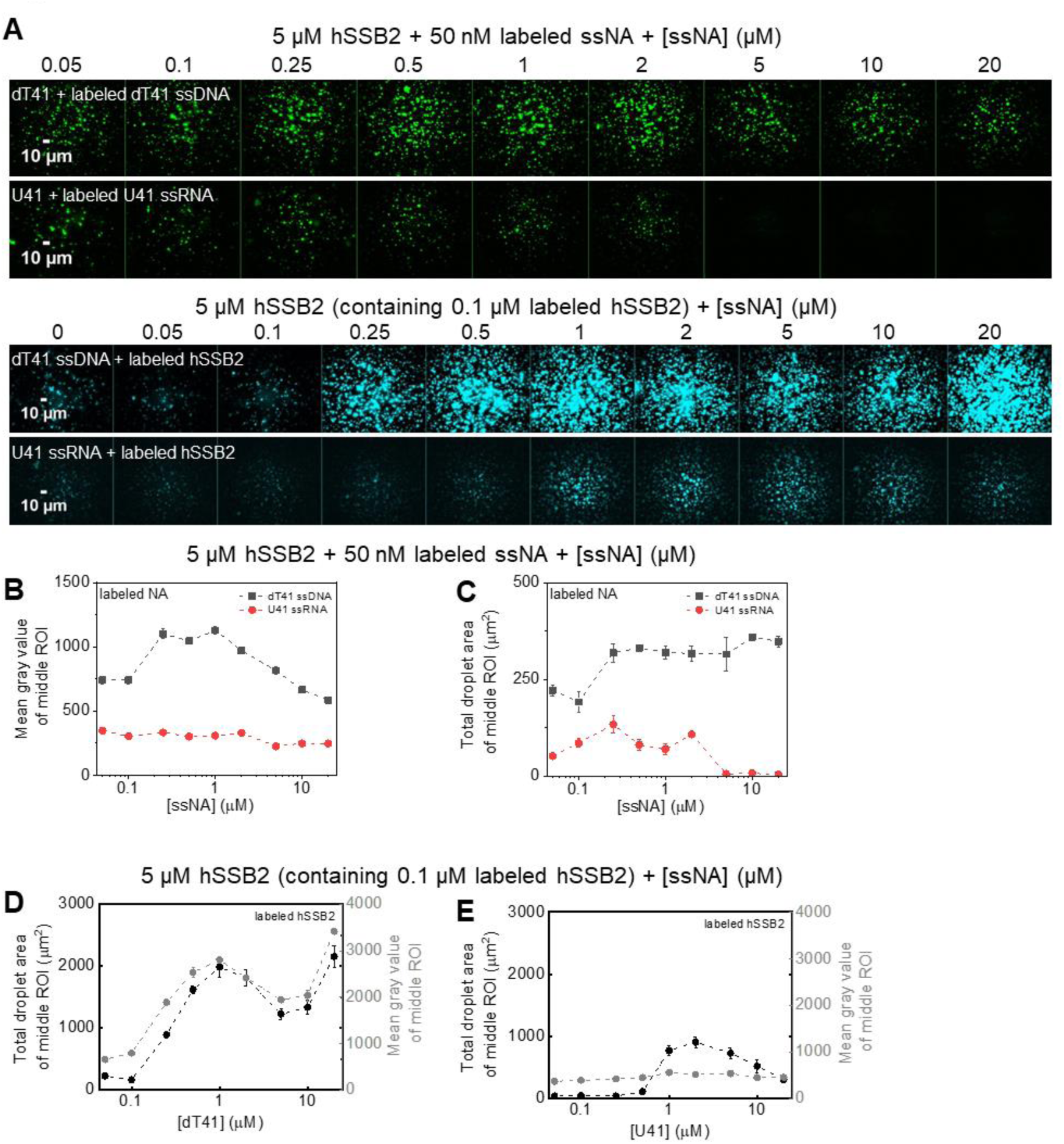
ssDNA induces hSSB2 condensation markedly more effectively than ssRNA. **(A)** Top rows: fluorescence microscopic images obtained upon mixing 5 µM hSSB2, 50 nM Cy3-labeled dT41 ssDNA or Cy3-labeled U41 ssRNA and indicated concentrations of unlabeled dT41 ssDNA or U41 ssRNA, respectively. Samples were incubated for 30 minutes before imaging. Bottom rows: dT41 and U41 concentration dependence monitored as above but in the presence of 0.1 µM fluorescein-labeled hsSSB2 instead of labeled oligos. Samples were incubated for 60 minutes before imaging **(B-C)** ssDNA (black) and ssRNA (red) concentration dependence of **(B)** mean gray values and **(C)** total droplet areas obtained from the center areas of fluorescence images shown in panel **A** top rows (see also Methods). Means ± SEM are shown for *n* = 3 **(D-E)** ssDNA and ssRNA concentration dependence of total droplet areas (black) and mean gray values (gray) determined from the center areas of fluorescence images in experiments performed with labeled hSSB2 (*cf.* panel **A** bottom rows). Means ± SEM are shown for *n* = 3.

In addition to the effect of nucleic acids, we also tested the contribution of the intrinsically disordered C-terminal region (IDR) of hSSB2 to LLPS (**Fig. 4**). Deletion of the IDR moderately decreased the protein’s affinity to ssDNA (**Fig. 4B, Table S2**). At the same time, IDR-deleted hSSB2 (hSSB2-dIDR) failed to form LLPS condensates (**Fig. 4C**) even under conditions that are quasi-saturating in terms of ssDNA binding (**Fig. 4B**). While hSSB2-dIDR solutions appeared transparent, at high hSSB2-dIDR concentrations we observed amorphous aggregate-like structures (**Fig. 4C**), similar to those seen for full-length hSSB2 in the absence of nucleic acids and in the presence of low amounts of ssRNA (cf. DIC results in **Fig. 2A** and U41 results in **Fig. 3A**).

**Figure 4.**
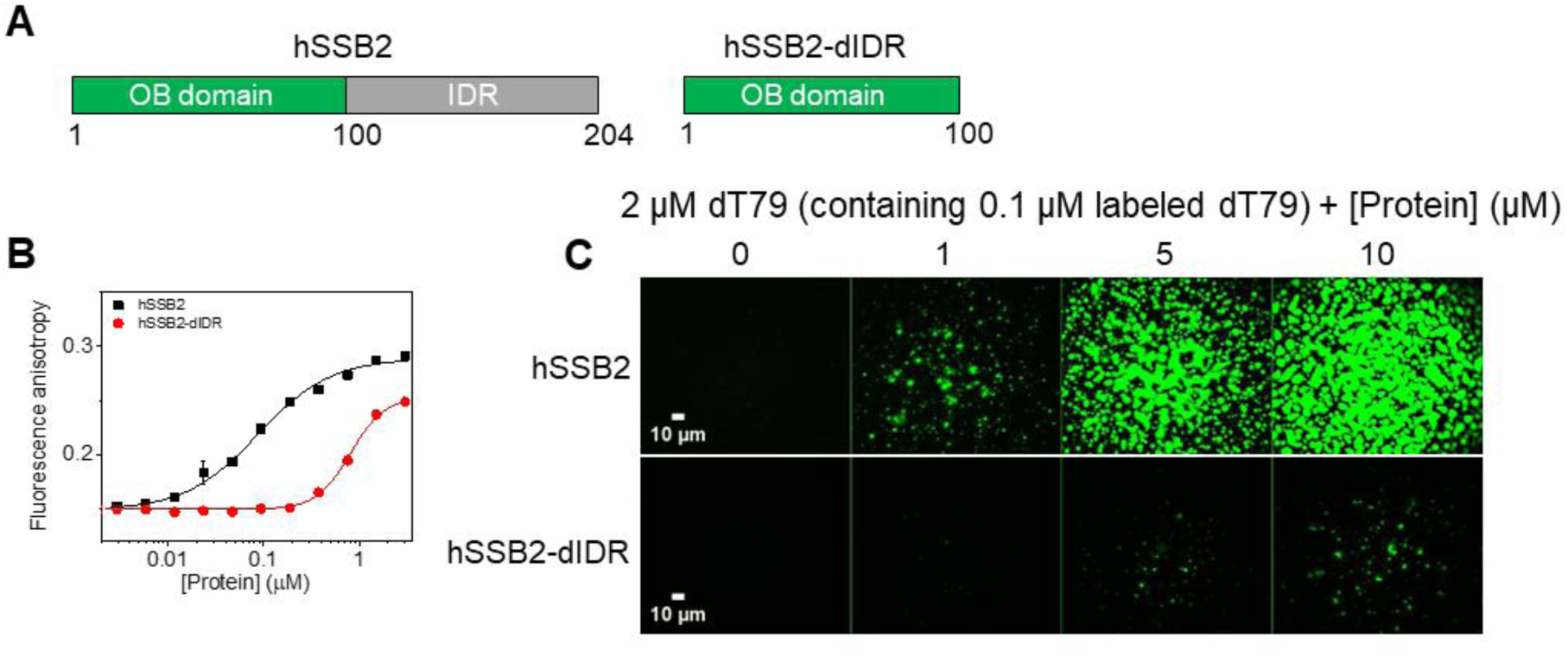
hSSB2 LLPS is mediated by the IDR region. **(A)** Schematic domain structure of full-length and IDR-truncated (dIDR) hSSB2 constructs. **(B)** Fluorescence anisotropy measurements performed by titrating 10 nM ss54 (3’- fluorescein labeled 54-mer ssDNA) with increasing concentrations of hSSB2 or hSSB2-dIDR. Solid lines show best fits using the Hill equation. Determined parameters are reported in **Supplementary Table 2**. Means ± SD are shown for *n* = 2. **(C)** Fluorescence microscopic images obtained upon mixing 2 µM dT79 (containing 0.1 µM Cy3-dT79) and increasing concentrations of hSSB2 or hSSB2-dIDR. Samples were mixed in a buffer containing 4 mM KCl and 8 mM MgCl_2_ instead of the standard 50 mM and 10 mM MgCl_2_ concentrations, and visualized after 30 minutes.

Taken together, the above results highlight that hSSB2 LLPS is more strongly enhanced by ssDNA than ssRNA, despite the protein’s ability to bind ssRNA; and that hSSB2 condensation is dominantly mediated by the IDR region.

### hSSB2 condensation occurs in a physiologically plausible, low protein concentration regime

Next, we aimed to determine how hSSB2 protein concentration influences LLPS by performing fluorescence microscopy experiments using fixed 2 µM dT79 concentration (containing 0.1 µM fluorescently labeled dT79) titrated with increasing hSSB2 concentrations (**Fig. 5A**). After mixing, condensed particles were detectable near the microscope slide surface, starting at 0.5 µM hSSB2 concentration (**Fig. 5A**), in line with rapid LLPS initiation observed in turbidity experiments (**Fig. 2F**). The number, size and intensity of observed particles increased with protein concentration, ultimately leading to saturation of the fluorescence signal (**Fig. 5B-C**). Both the number and size of detected particles increased with increasing incubation time and, after 1 h, particles were observed even at 0.1 µM protein concentration (**Fig. 5**). The time-dependent appearance of observable hSSB2 droplets may result either from *de novo* droplet formation and growth, or from fusion of smaller particles that are below the detection limit. As the N-terminal histidine tag present in our constructs may influence LLPS characteristics, we also purified tag-free hSSB2 and compared its protein concentration dependent condensation with that of His-tagged hSSB2 (**Fig. S6A**). Results indicated that the presence of the histidine tag moderately enhances condensation: condensates were observed at 5 µM and higher protein concentrations for the tag-free variant (**Fig. S6B**). At higher protein concentrations, both constructs behaved similarly (**Fig. S6B-C**). Turbidity measurements confirmed that the histidine tag has a moderate enhancing effect on hSSB2 LLPS at lower protein concentrations (**Fig. S6D**). In summary, these results indicate that microscopically observable hSSB2 phase separation can readily occur in the low µM protein regime.

**Figure 5.**
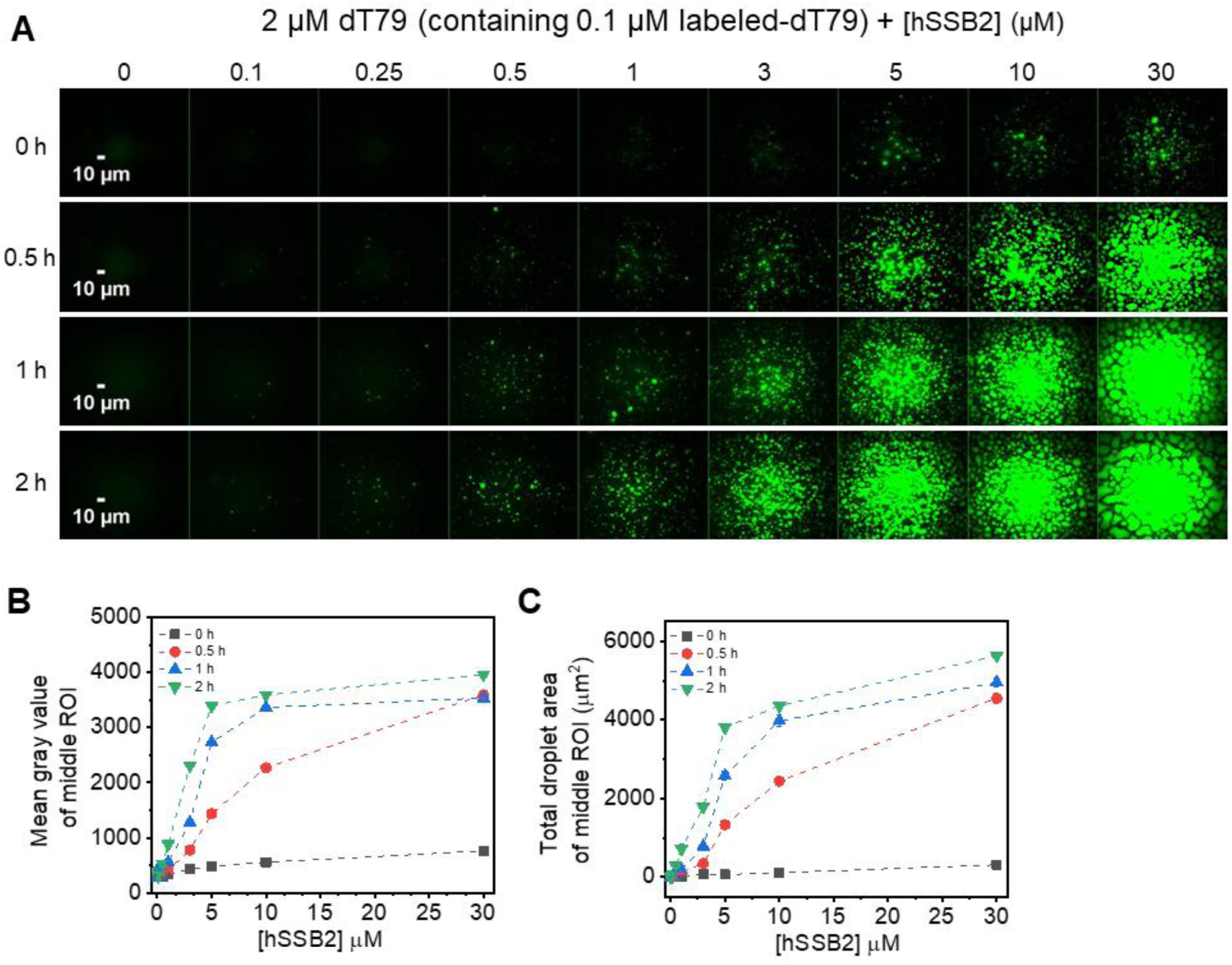
Robust hSSB2 phase separation occurs in the low micromolar protein concentration regime. **(A)** Fluorescence microscopic images obtained upon mixing 2 µM dT79 (containing 0.1 µM Cy3-dT79) and indicated concentrations of hSSB2. Samples were incubated in standard LLPS buffer for the indicated times **(B-C)** hSSB2 concentration dependence of **(B)** mean gray values and **(C)** total droplet areas obtained from the center areas of fluorescence images shown in panel **A**. Means ± SEM are shown for *n* = 3.

### hSSB2 condensation is sensitive to the redox environment and occurs under physiological ionic conditions

hSSB2 and hSSB1 contain three conserved cysteine residues (two in the OB domain (C45 and C85 in hSSB2) and one (C103) at the base of the IDR) that were shown to facilitate the formation of oligomers held together by disulfide bonds in case of hSSB1 (28). This finding raises the possibility that the redox state of hSSB2 may influence LLPS characteristics. To test this idea, we performed fluorescence microscopy experiments in the presence of increasing concentrations of reducing (DTT) and oxidizing (H_2_O_2_) agents (**Fig. 6A-B**). In the absence of ssDNA, we observed only a small number of aggregate-like structures in all conditions tested (**Fig. 6A**). However, in the presence of ssDNA, redox conditions markedly influenced the LLPS behavior (**Fig. 6B-D**). Under reducing conditions, condensates appeared more rapidly and in larger numbers, were larger in size and in general had more droplet-like appearance compared to particles observed under oxidizing conditions. Even low concentrations of H_2_O_2_ were sufficient to alter LLPS characteristics (**Fig. 6B-D**). We also tested the effect of redox conditions and DTT removal in experiments using higher hSSB2 concentration (10 µM). Both DTT-free (dialyzed) and DTT-containing samples showed limited aggregation in the absence of nucleic acids, and condensation was apparent upon ssDNA addition in all cases (**Fig. 6E-G**). However, in the DTT-removed sample (**Fig. 6E**) the number and morphology of observed particles were markedly different from those in 1 or 50 mM DTT-containing controls (**Fig. 6F-G**). The effect was especially prominent at longer incubation times, as in the absence of DTT the protein formed amorphous structures resembling branched strings of spheres. Using the same protein concentration, similar structures were also observed upon longer incubation in the presence of H_2_O_2_, whereas condensates incubated with TCEP reducing agent retained their disperse, spherical, droplet-like form even for 24 hours (**Fig. 6H**). Branched structures observed in H_2_O_2_ were stable without showing signs of fusion and, based on the observed high fluorescence signal, hSSB2 likely retained its ssDNA binding ability. In summary, the above experiments revealed that hSSB2 undergoes *bona fide* LLPS under reducing conditions, whereas oxidative conditions promote formation of solid-like nucleoprotein particles.

**Figure 6.**
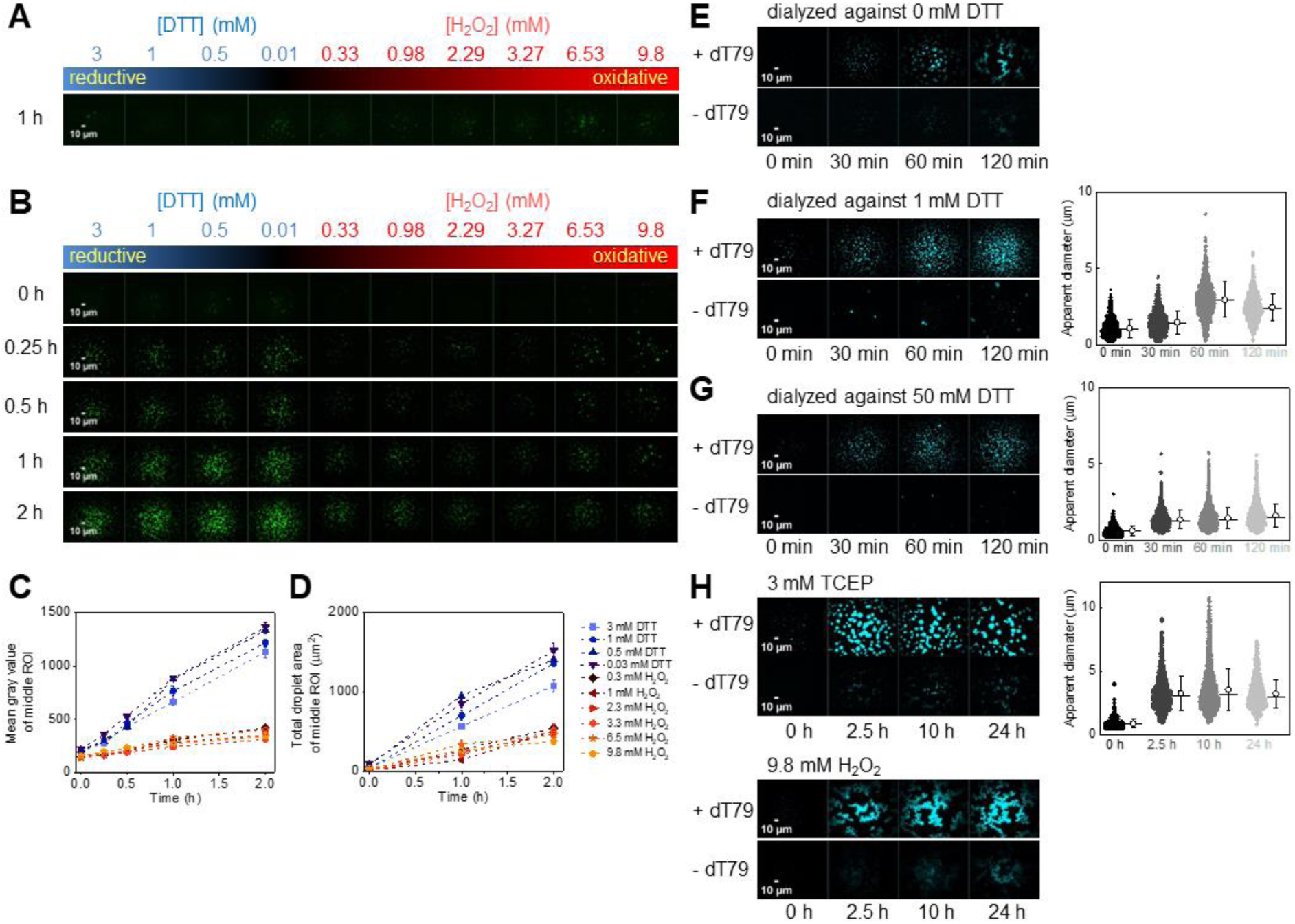
hSSB2 forms liquid-like and solid-like condensates under reducing and oxidizing conditions, respectively. **(A-B)** Fluorescence microscopic images obtained upon mixing (**A**) 4.9 µM hSSB2 with 0.1 µM fluorescein-labeled hSSB2, or (**B**) 5 µM hSSB2 and 2 µM dT79 containing 0.1 µM Cy3-dT79 in standard LLPS buffer containing the indicated DTT (reducing agent) or H_2_O_2_ (oxidizing agent) concentrations. Samples in panel **A** were incubated for 30 min before imaging. Incubation times for panel **B** are indicated in the figure **(C-D)** DTT and H_2_O_2_ concentration and incubation time dependence of **(C)** mean gray values and **(D)** total droplet areas obtained from the center areas of fluorescence images shown in panel **B**. Means ± SEM are shown for *n* = 3. **(E-G)** Left: Fluorescence microscopic images obtained after incubating 10 µM hSSB2 in the absence of DTT **(E)**, in 1 mM DTT **(F)** or 50 mM DTT **(G)** with 0.1 μM fluorescein-labeled hSSB2 in the presence or absence of 1 μM dT79 for the indicated durations. Buffers contained DTT at concentrations identical to those in the corresponding hSSB2 stock solutions. Right: particle size distributions determined from microscopic images. Calculated means (circles) ± SD and medians (horizontal lines) are shown to the right of each distribution. **(H)** Fluorescence microscopic images obtained upon incubating 10 μM hSSB2 (containing 0.1 μM fluorescein-labeled hSSB2) with and without 1 μM dT79 in standard LLPS buffer containing 3 mM TCEP reducing agent or 9.8 mM H_2_O_2_ for the indicated durations. Particle diameter distributions and calculated means (circles) ± SD and medians (horizontal lines) obtained in the presence of TCEP are shown on the right.

LLPS systems are often sensitive to the ionic milieu (41,42). Accordingly, during protein purification we observed that the addition of MgCl_2_ decreases the turbidity of highly concentrated hSSB2 solutions. To elucidate the effect of ionic conditions, we monitored hSSB2 condensation in the presence of ssDNA and various salt types and concentrations (**Fig. 7**). MgCl_2_ and KCl inhibited condensation in a concentration-dependent manner based on fluorescence microscopy (**Fig. 7A-E**) and turbidity results (**Fig. 7F**). Control measurements confirmed that the effect is not influenced by the presence of the histidine tag (**Fig. S7**). Complete LLPS inhibition was observed at about 2 times lower MgCl_2_ concentrations than that for KCl, indicating that Cl^-^ ions dominantly contribute to the inhibitory effect (**Fig. 7C-D, F**). In line with this observation, NaCl showed similar effect to that of KCl (**Fig. 7E-F**). Na- glutamate and K-glutamate did not fully inhibit condensation even at high concentrations; however, the turbidity of the solution moderately decreased with increasing concentrations of glutamate salts (**Fig. 7F**). Additional microscopy experiments performed at a fixed 250 mM concentration of various salts corroborated the above results and also revealed that CaCl_2_ shows effects similar to those of MgCl_2_, whereas the effect of Mg-acetate is similar to those of KCl and NaCl (**Fig. 7E**). Taken together, Cl^-^ and acetate^-^ have a similar inhibitory effect on hSSB2 LLPS, whereas glutamate is permissive. Interestingly, neither the type nor the concentration of cations influence hSSB2 LLPS, as CaCl_2_ and MgCl_2_ behaved similarly; the effects of KCl, NaCl and Mg-acetate were similar; and both Na- and K-glutamate were permissive. Additionally, we observed that Cl^-^ containing salts (KCl and MgCl_2_) practically eliminated hSSB2 binding to ssDNA at high concentrations (250 mM). In contrast, K- glutamate had a less pronounced effect, reducing the binding affinity by ∼6-7 fold without altering cooperative binding, consistent with its milder impact on LLPS (**Fig. 7G**, cf. **Fig. 1C**). These findings collectively suggest that salt dependent LLPS inhibition occurs through the attenuation of binding to ssDNA, with certain anions, particularly Cl^-^ and acetate, exerting the strongest effect. Importantly, however, the ion concentrations required for complete LLPS inhibition (∼500 mM Cl^-^, >250 mM acetate) are markedly higher than the respective physiological concentrations (10 - 100 mM Cl^-^, < 1 mM acetate, 100 mM K^+^, 10 mM Na^+^, 10 mM Mg^2+^, 0.1 – 1 µM Ca^2+^, < 20 mM glutamate (43–45)), indicating that the intracellular ionic milieu is permissive for hSSB2 LLPS.

**Figure 7.**
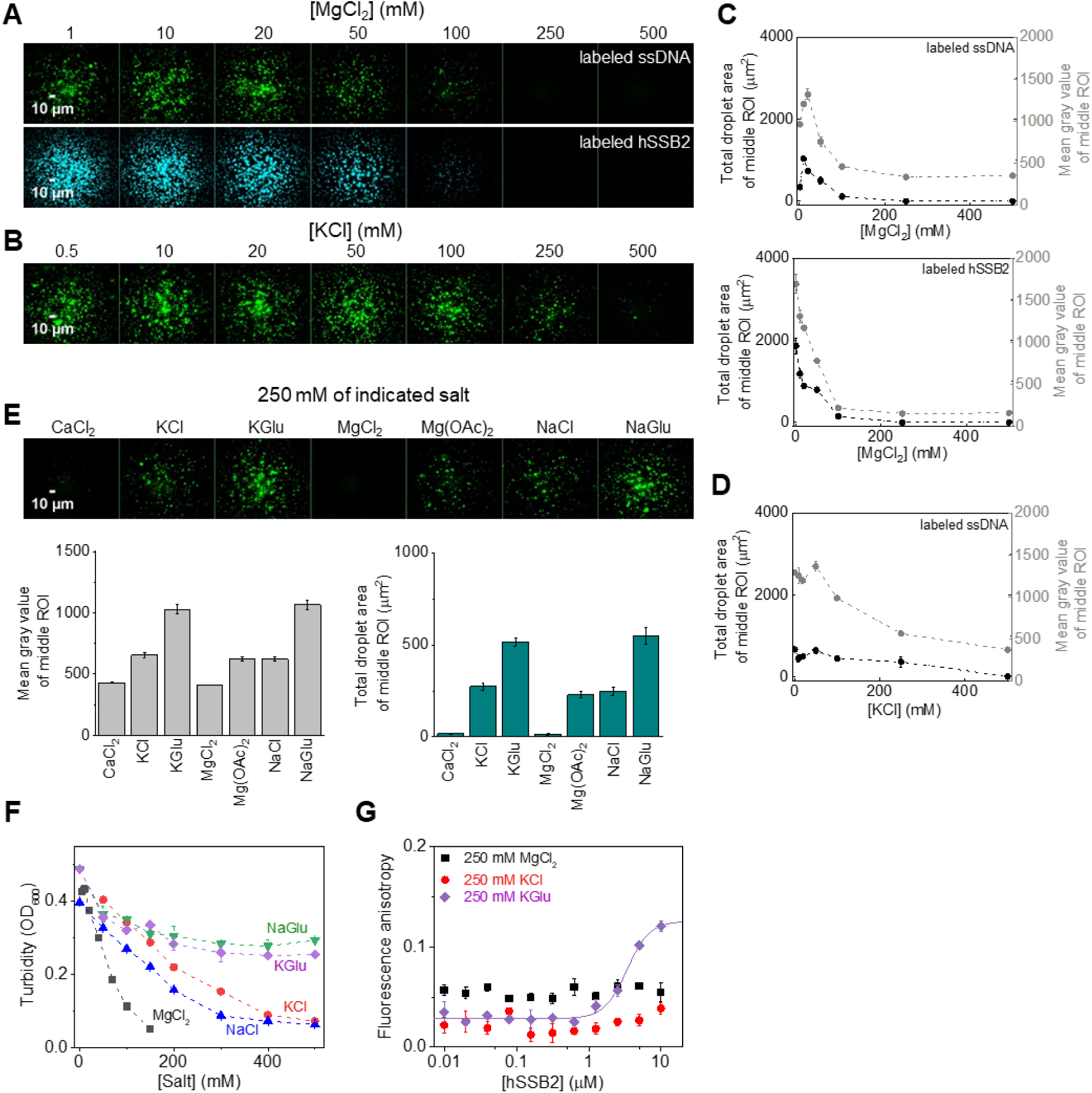
hSSB2 condensation occurs in cellular-like ionic conditions. **(A)** Fluorescence microscopic images of samples obtained after incubating (top row) 5 µM hSSB2 and 2 µM dT79 containing 0.1 µM Cy3-labeled dT79, or (bottom row) 10 µM hSSB2 containing 0.1 µM fluorescein-labeled hSSB2 and 2 µM dT79 for 30 and 60 minutes in LLPS buffer containing indicated concentrations of MgCl_2_ and 50 mM KCl. **(B)** Fluorescence microscopic images obtained as in the top row of panel **A**, but in the presence of indicated KCl concentrations and 10 mM MgCl_2_ **(C-D)** Salt concentration dependence of total droplet area and mean gray value of middle ROIs determined from experiments shown in panels **A-B**. Means ± SEM are shown for *n* = 3. **(E)** Top row: fluorescence microscopic images obtained after 30-min incubation of 5 µM hSSB2 and 2 µM dT79 (containing 0.1 µM Cy3-labeled dT79) in LLPS buffer containing only the indicated salts at 250 mM concentration. Bottom row: salt type dependence of the total droplet area and mean gray value of middle ROIs. Means ± SEM are shown for *n* = 3. **(F)** Salt type and concentration dependence of the turbidity of solutions containing 20 µM hSSB2 and 2 µM dT79. Samples were incubated in LLPS buffer containing the indicated concentrations of salts, and measured after 5 minutes. MgCl_2_ titrations contained additional 50 mM KCl. All other measurements contained 10 mM MgCl_2_. Means ± SEM are shown for *n* = 3. **(G)** Fluorescence anisotropy measurements performed by titrating 10 nM 3’-fluorescein labeled, 36-mer ssDNA (ss36, cf. Fig. 1C) with increasing concentrations of hSSB2 in the presence of 250 mM KCl or MgCl_2_ or K-glutamate (KGlu). Solid line shows best fit to K-glutamate data using the Hill equation (*K*_d_ = 3.4 ± 0.3 μM, *n* (Hill coefficient) = 2.8 ± 0.6). Means ± SEM are shown for *n* = 3.

### Protein interaction partners of hSSB2 selectively enter and become enriched in hSSB2 condensates

hSSBs were previously shown to interact with a multitude of proteins associated with various cellular processes (13,19). Therefore, we tested whether previously identified hSSB2 interaction partners, INTS3 and INIP (13) can partition into hSSB2 nucleoprotein coacervates, using a dual detection-based fluorescence microscopic assay and fluorescently labeled variants of the investigated partners (**Fig. 8A**). We also tested whether the paralogous hSSB1 protein can be enriched inside hSSB2 condensates. As hSSB1 was shown to interact with Bloom’s syndrome (BLM) helicase (20), we also tested the partitioning of BLM and, separately, that of the Topoisomerase IIIa-RMI1-RMI2 (TRR) complex that functions together with BLM during DNA repair processes (46–49). In addition to the fluorescently labeled partners, hSSB2 nucleoprotein coacervates were directly visualized using fluorescently labeled ssDNA. In the absence of hSSB2, none of the tested proteins showed condensation at the applied 0.1 µM concentration, and only a diffuse ssDNA signal was observed (**Fig. 8B**). However, all indicated interaction partners and, strikingly, hSSB1, BLM and the TRR complex showed enriched partitioning in hSSB2 condensates (**Fig. 8B-C**). Importantly, INTS3 and INIP bind ssDNA only weakly (12), implying that their partitioning into hSSB2 condensates is driven by protein-protein rather than protein-DNA interactions. Interestingly, INTS3 colocalized to a lesser extent with hSSB2- ssDNA coacervates than did the other protein partners (**Fig. 8C**). This effect may stem from the higher noise present in the INST3 signal or from compositional heterogeneity of condensates in the presence of INTS3. In contrast to hSSB2’s interaction partners, neither enhanced green fluorescent protein (EGFP) nor EcSSB showed enrichment in hSSB2 droplets, reflecting the action of hSSB2 condensates as molecular filters allowing selective enrichment based on specific molecular interactions (**Fig. 8B-C**). In line with this, small molecules (fluorescein, fluorescein-deoxycytidine (dCTP)) or fluorescently labeled peptides (ones comprising the last 8 amino acids of EcSSB and hSSB1), which likely cannot interact specifically with hSSB2, showed only weak to moderate signal correlation to the hSSB2 coacervate signal without enrichment in hSSB2 coacervates (**Fig. 8D-E**). These results indicate that while hSSB2 coacervates are permeable for both macromolecules and small molecules (exclusion was not observed in any case), enrichment requires specific interactions with hSSB2, and ssDNA binding by another protein is not sufficient for co-condensation.

**Figure 8.**
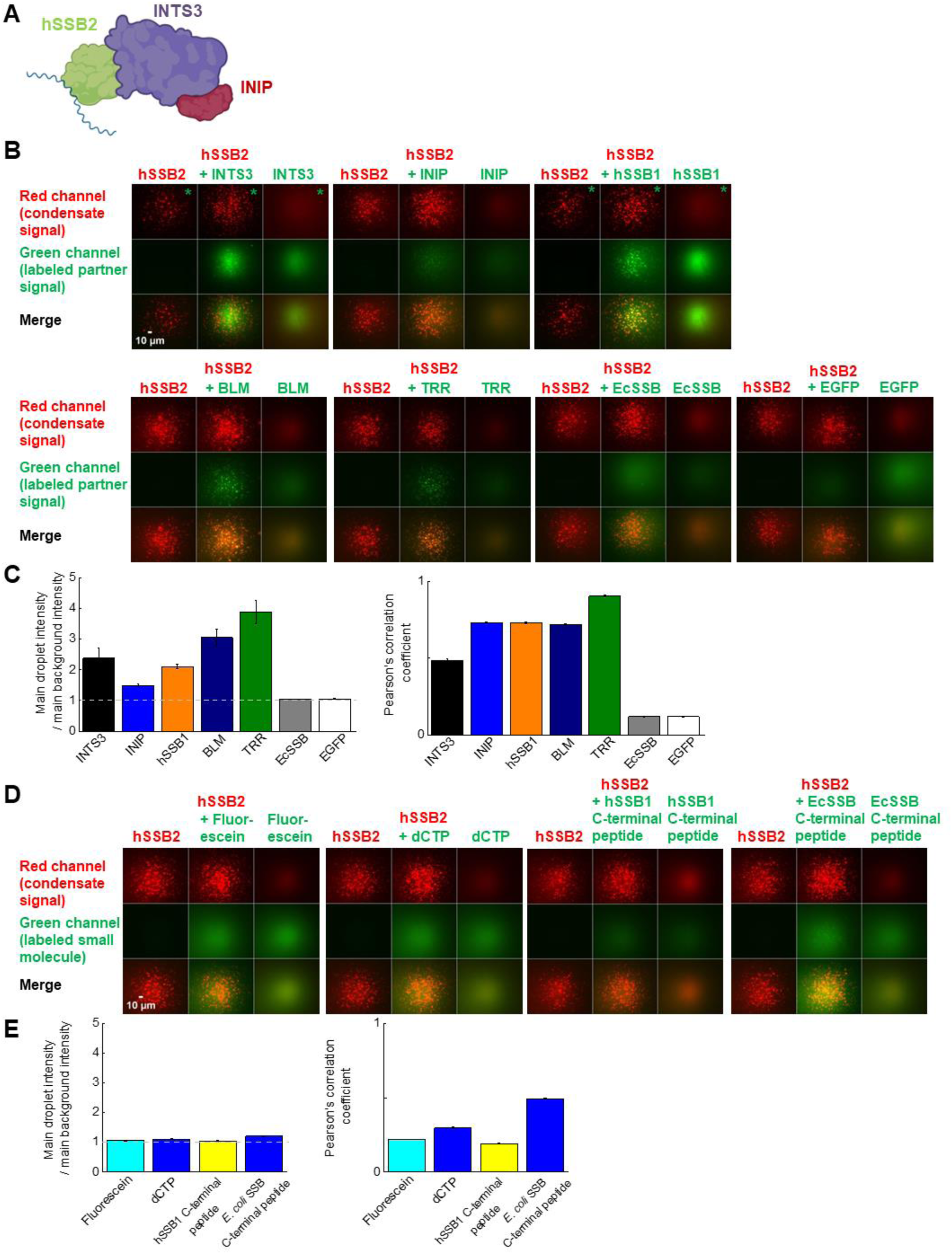
Protein interaction partners of hSSB2 selectively partition into hSSB2 condensates. **(A)** Schematic image of the SOSS (Sensor of ssDNA) complex comprised of the INTS3 and INIP proteins, and hSSB2 bound to ssDNA (blue line). **(B)** Two-laser fluorescence microscopic images obtained from experiments monitoring the co-condensation of hSSB2 and investigated protein partners (INTS3, INIP, hSSB1, human BLM helicase (BLM), Topoisomerease IIIa-RMI1-RMI2 (TRR) complex, *E. coli* SSB, or enhanced green fluorescent protein (EGFP)). For each tested partner, columns represent 3 parallel experiments in which either 5 µM hSSB2, 5 µM hSSB2 plus 0.18 µM labeled protein partner, or 0.18 µM labeled partner alone were incubated with 2 µM dT79 containing 0.1 µM fluorescently labeled ssDNA. Condensates were visualized via the ssDNA signal. Fluorescent probe pairs (listed in Methods) were chosen to avoid cross-detection of ssDNA and investigated protein partner signals (arbitrary colored red and green, respectively). The bottom panel of each column shows the merge of the simultaneously monitored two signals. Images were not background corrected. **(C)** Left: Enrichment of protein partners in hSSB2 condensates, calculated as the ratio of the mean protein partner signal intensity localized inside hSSB2 droplets and the mean background intensity. Right: Pearson’s correlation coefficients calculated from the hSSB2 droplet signal (monitored via labeled ssdT79) and indicated partner signals from dual-channel fluorescence microscopic images (see middle columns in each tile in panel **B**). Means ± SEM are shown for *n* = 3. **(D-E)** Two-laser fluorescence microscopic experiments monitoring co-condensation of fluorescein, fluorescently labeled deoxycytidine (dCTP), or fluorescently labeled peptides (last 8 amino acids of EcSSB and hSSB1) with hSSB2. Experiments were performed as described for protein partners in panel **A** and analyzed as in panel **C**. Means ± SEM are shown for *n* = 3.

## DISCUSSION

In this study we show that hSSB2 can undergo liquid-liquid phase separation (**Fig. 2**) mediated by its intrinsically disordered C-terminal region (**Fig. 4**). The process requires the presence of ssDNA, and is facilitated by ssRNA to a much more limited extent (**Figs. 2, 3, S5**), despite the protein’s similar binding affinity to both types of nucleic acid (**Fig. 1C, S2**). The LLPS-enhancing effect depends on ssDNA length and also on hSSB2:ssDNA stoichiometry (**Fig. 2B-D**, F-J, **S5**). We found that hSSB2 condensation is triggered by ssDNA molecules of 15 or more nucleotides, and requires at least 32 nucleotides for maximal stimulation (**Fig. 2G-J**). hSSB2 and hSSB1 were proposed to exist dominantly as monomers in solution; however, hSSB2 multimerization was observed upon longer storage of preparations and was proposed to occur also upon ssDNA binding (10,27). The structure of hSSB2 has not been determined experimentally in any state; however, the hSSB1 monomer interacts with ∼7 nucleotides of ssDNA in available crystal structures, and a similar ssDNA binding mode was proposed for hSSB2 monomers (50). Indeed, previous experimental results indicated that hSSB2 can interact with such short DNA molecules, but the binding affinity was shown to increase with ssDNA length, and 15 - 30 and 20 - 30 nt binding site size ranges were estimated for mouse SSB2 and hSSB2, respectively (2,10). In our experiments, dT7 was unable to stimulate hSSB2 condensation, possibly due to its low affinity to hSSB2 (**Fig. 2G**). Notably, we observed a similar ssDNA length dependent effect on hSSB2 condensation stimulation as described previously for binding affinity, and the proposed binding site size range aligns well with our determined ssDNA length (∼32 nt, **Fig. 2H**) required for maximal condensation stimulation. It must be noted that the mechanistic interpretation of the above discussed binding site and optimal condensation-stimulating ssDNA length values depends on the oligomerization state and number of SSB2 monomers in contact with ssDNA. Our results suggest that the intrinsic binding site size of a hSSB2 monomer may be lower or near the lower limit of the proposed binding site size range, and ssDNA molecules that are longer than this size may either act as a scaffold to which multiple hSSB2 molecules bind, or ssDNA binding may induce hSSB2 oligomerization. Accordingly, we observed cooperative binding with 36-mer and 41-mer nucleic acids and the presence of multiple complex forms in EMSA experiments. Furthermore, LLPS reached maximal efficiency at substoichiometric ssDNA concentrations for lengths greater than 15 nt, whereas high ssDNA concentrations decreased condensation efficiency (**Fig. 2B, G, 3A, Fig. S2**). While we cannot at this point distinguish between various binding and oligomerization scenarios, in either case ssDNA binding can be stabilized by intermolecular hSSB2 interactions, could increase the protein’s local concentration, and bring about multivalency for intermonomer IDR-IDR interactions, thereby facilitating condensation. Based on the proposed scaffolding effect, high ssDNA concentrations likely inhibit LLPS by decreasing the probability of multiple hSSB2 molecules binding to the same ssDNA strand and/or due to competition of excess nucleic acids for surfaces on hSSB2 that could drive LLPS *via* protein-protein interactions. However, ssDNA apparently has an additional, nucleic acid type-specific effect, as ssRNA only weakly induced hSSB2 condensation (**Fig. 3, S5**), despite the interaction of multiple hSSB2 molecules with the tested RNA molecules, as indicated by the cooperative ssRNA binding behavior and the presence of multiple complex forms in electrophoretic mobility shift assays (**Fig. 1C, S2**). A possible explanation for these findings is that ssDNA binding specifically elicits a nucleoprotein configuration (and/or hSSB2 conformation) that is more prone to condensation than that adopted upon ssRNA binding. Further experiments will be required the explore the molecular details underlying ssDNA-specific condensation and the oligomerization state of hSSB2 upon nucleic acid binding.

We also observed that the redox environment influences hSSB2’s phase separation propensity (**Fig. 6**). Oxidation apparently decreases the fluidity of coacervates and induces a more solid-like state, suggested by the formation of filamentous, branched structures comprised of bead-like particles and the lack of fusion between particles, in contrast to singular, droplet-like structures and their fusion observed under reducing conditions. As inferred from the crystal structure of hSSB1, hSSB2 may harbor two surface-exposed cysteines that could mediate covalent oligomerization by disulfide bridge formation (28,50), as well as other surface exposed amino acids that can undergo oxidation (methionine, tyrosine, tryptophan, histidine). Mutational studies on cysteine-to-serine substituted hSSB1 variants highlighted the role of C41 and C99 (homologous to C45 and C103 of hSSB2, respectively) in oxidation-induced covalent dimerization (28,39), and the presence of all three hSSB1 cysteines (C41, C81, C99) was shown to be required for redox-dependent LLPS (albeit not a prerequisite for LLPS *per se*) (39). Whether oxidation alters LLPS characteristics of hSSB2 by inducing covalently linked forms or *via* other oxidation-mediated structural changes remains a question for further studies. While the structural basis of ssDNA binding by hSSB1 and hSSB2 OB folds appears similar, it has been proposed that the divergent IDRs of the two paralogs may drive differences protein-protein interaction properties (27). Similarly, the divergent redox dependent LLPS properties of hSSB1 and hSSB2 are likely to be determined in large part by the divergent IDRs of the two proteins.

The available body of observations on hSSB2 raises the important question whether hSSB2 condensation occurs in cells and if so, what kind of functional advantage could be conferred by such a mechanism. Physiological ionic conditions apparently permit hSSB2 LLPS (**Fig. 7**). We found condensation to be specifically inhibited by Cl^-^ (and acetate), whereas mono- and divalent cations or glutamate had little effect. DNA binding measurements revealed that the DNA binding affinity of hSSB2 is sensitive to the mentioned salts. Specifically, a high concentration of potassium glutamate led to a decrease in binding affinity by a factor of ∼6-7, while cooperative binding remained unaffected (**Figs. 1C** and **7G**). In contrast, salts containing chloride ions (KCl, MgCl_2_) drastically weakened ssDNA binding at the same concentration (**Fig. 7G**). These results suggest that Cl^-^ (and acetate) specific LLPS inhibition is primary due to attenuated ssDNA binding. Interestingly, Cl^-^ showed a similar LLPS inhibitory effect in case of EcSSB, whereas glutamate enhanced condensation of that SSB homolog in the absence of nucleic acids (32,51). This raises the possibility that intermolecular hSSB2 interactions, involved in the cooperative binding mechanism that significantly increases ssDNA binding affinity and are essential for LLPS, may also be influenced by the type and concentration of salt. Hence, exploring the detailed mechanism of ion-dependent effects will be important in future studies. Importantly, all observed LLPS inhibitory ion concentrations were markedly higher than their proposed intracellular concentrations. Besides the ionic milieu, LLPS also depends on protein concentration (**Fig. 5, S6**). The cellular concentration of hSSB2 has not been investigated in detail, although a high-throughput mass spectroscopy study estimated a copy number of about 5,000 hSSB2 molecules, in contrast to about 500,000 hSSB1 molecules, in HeLa cells (52). Assuming a dominantly nuclear localization and a 374 µm^3^ nuclear volume for HeLa cells (53), these values translate to 0.02 and 2.22 µM for hSSB2 and hSSB1, respectively. In line with this difference, the mRNA level of hSSB1 was found to be significantly higher in HeLa cells than that for hSSB2 (68.2 nTPM versus 11.7 nTPM, respectively; Human Protein Atlas database, cell line data). Importantly, however, hSSB2 expression is tissue specific, with epithelial and immune cells showing high transcription levels (the highest level, 345.3 nTPM, was detected in Langerhans cells (Human Protein Atlas database, single-cell RNA data)). Moreover, hSSB2 expression may well be enhanced upon genomic stress, as indicated by the protein’s involvement in genome maintenance processes and its increased expression levels upon hSSB1 ablation or UV-stress (5,14). All in all, hSSB2 levels may greatly vary among cell types, and robust LLPS may manifest in cells showing high expression levels. Microscopically observable protein droplets readily formed in the low micromolar concentration regime (**Fig. 5, S6**) *in vitro* even in the absence of molecular crowders (e.g. proteins) that may further enhance condensation in the cell (54). In addition, hSSB2 binding to longer ssDNA stretches may greatly increase the local protein concentration and thus could potentiate IDR-IDR interactions and condensation even at low cellular hSSB2 levels. hSSB1 and hSSB2 co-condensation, possibly in conjunction with SOSS complex components, is another plausible mechanistic scenario.

In line with the above proposition, we recently discovered that, besides hSSB2, hSSB1 is also prone to undergo phase separation (39). However, the LLPS propensity of the two human SSBs differs in several important aspects. Although phase separation of both SSBs is induced by nucleic acids, hSSB1 also requires an oxidative environment for LLPS. The presence of the IDR is essential for LLPS in both proteins. Intriguingly, besides ssDNA-induced LLPS, ssRNA also strongly induced condensation by hSSB1, in contrast to weak RNA induction of hSSB2 LLPS. Genome maintenance proteins that bind to hSSBs are selectively enriched in both hSSB1 and hSSB2 coacervates, while non-interacting proteins are excluded. Furthermore, each of the two SSBs is able to enter condensates formed by the other paralog, suggesting the potential existence of mixed hSSB1-hSSB2 condensates in cells during genome maintenance processes.

As another notable implication, our data reveal a markedly different linkage between nucleic acid binding and LLPS in studied bacterial *versus* eukaryotic SSB homologs, which might signify a functional adaptation to spatial relations within different cells (**Fig. 9**). We showed recently that EcSSB condensation is favored in the absence of nucleic acids, and ssDNA binding dissolves EcSSB condensates (32). In contrast, here we show that hSSB2 condensation requires ssDNA binding, similarly to that found recently for human RPA (38) and hSSB1 (39). In *E. coli*, ssDNA-induced solubilization of stored SSB particles can lead to rapid mobilization of SSB (and SSB-interacting protein) pools from condensates (∼100-nm diameter, ∼2,000 SSB tetramers) upon genomic stress. However, the more extended space and higher amount of material in the eukaryotic nucleoplasm would not allow for such rapid mobilization: for micron-sized particles, such mobilization could take minutes (cf. FRAP results on EcSSB diffusion (32)). Therefore, under these circumstances, it may be more adaptive to keep SSB molecules evenly dispersed in the nucleoplasm so that they are available at the sites of action, *i.e.*, sites of genomic damage (**Fig. 9**). This proposition is supported by our observation that hSSB2 is prone to undergo LLPS upon binding ssDNA molecules with a length of 32 or more nucleotide units. Although the subcellular localization of hSSB2 is yet unexplored, the above idea is further supported by the fact that hSSB1, presumably participating in the oxidative stress response, shows a diffuse nuclear and cytoplasmic localization in HeLa cells under normal conditions, but cytoplasmic granulation occurs when oxidative stress is applied (39).

**Figure 9.**
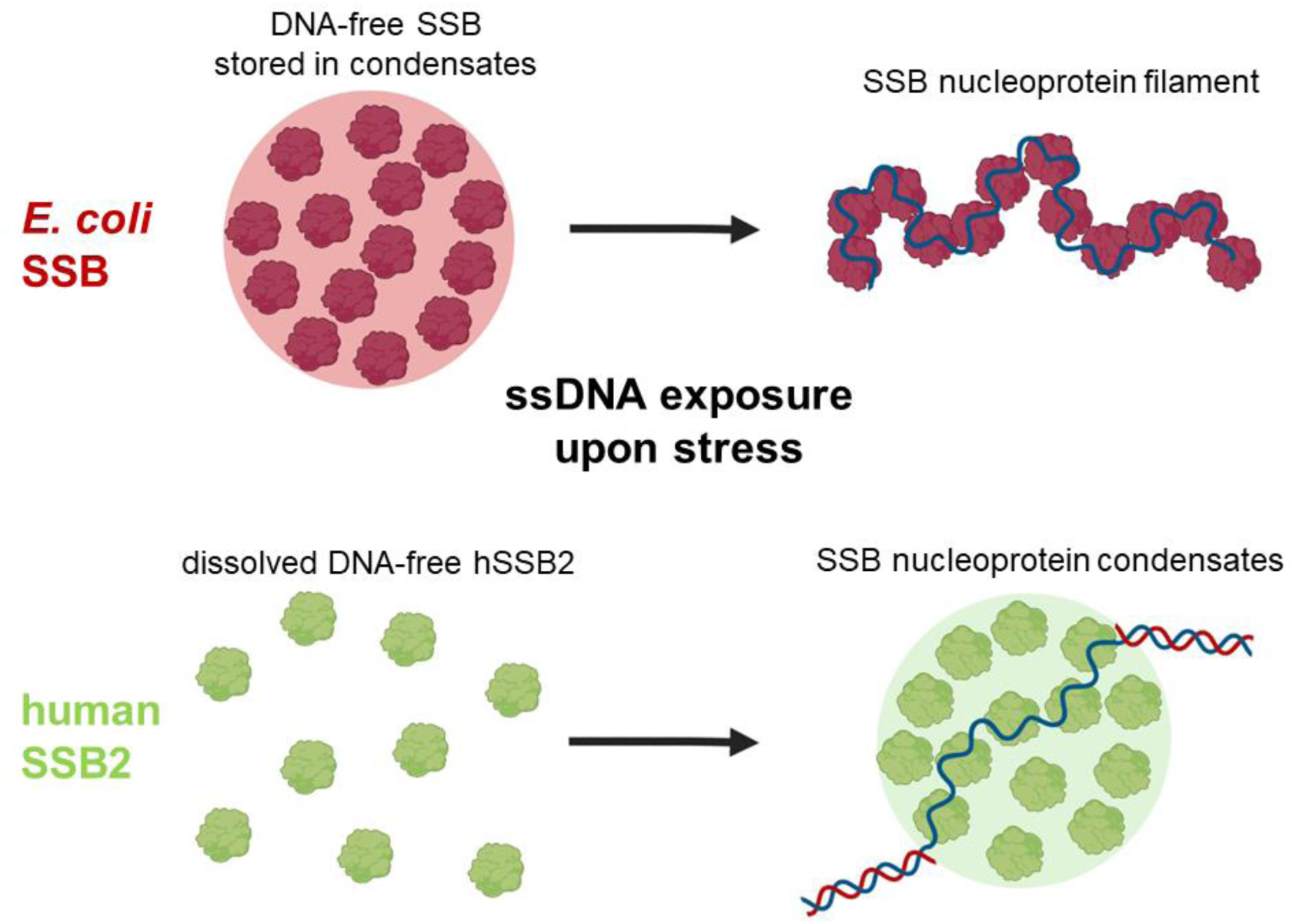
Proposed model for differential spatiotemporal regulation of SSB-driven LLPS in bacterial and eukaryotic cells. *E. coli* SSB forms condensates in an ssDNA-free form (32,51); thus, in the absence of genome stress, the majority of the cellular SSB pool is stored in membrane-associated LLPS foci(32,37). Upon exposure of excess ssDNA upon genome stress, solubilized SSB rapidly relocalizes to damaged genomic sites to form nucleoprotein filaments. In contrast, ssDNA-free hSSB2 is stored in soluble form in the nucleoplasm in absence of stress. hSSB2 condensates form upon binding to ssDNA stretches that arise during genomic stress.

Condensation induced by ssDNA exposure upon genome stress could facilitate formation of DNA repair compartments on damaged DNA regions. Supporting this proposition, we found that known (INTS3, INIP) and possible (hSSB1, BLM, TRR complex) hSSB2 interacting proteins partitioned into hSSB2-ssDNA coacervates (**Fig. 8**). At the same token, the low physiological hSSB2 expression levels observed in many tissues (Human Protein Atlas database) could be an adaptation to avoid ectopic LLPS and granule formation. hSSB2 overexpression may thus be detrimental for DNA repair, at least in part via uncontrolled LLPS. Furthermore, the differential expression levels and tissue patterns of hSSB2 and hSSB1 may be associated with the observed characteristic differences in their LLPS properties.

Generally high basal expression levels of hSSB1 could be adaptive in terms of oxidation-induced hSSB1 phase separation, whereas hSSB2 LLPS could be regulated by inducible protein abundance. hSSB2 appears to be dispensable for DNA replication and cell cycle progression under normal conditions, but it has important DNA repair functions under stress, and may thus be implicated as a useful target for cancer therapy (13–15,23,25,55,56). hSSB2 downregulation sensitizes cells for DNA damage, as evidenced by increased UV sensitivity, increased amounts cyclobutane pyrimidine dimer lesions, impaired RPA localization to DNA damage sites, and delayed recruitment of the XPC nucleotide excision repair protein upon UV irradiation (14,27). Furthermore, high hSSB2 expression was found to be associated with worse breast cancer prognosis, while downregulation of hSSB2 expression in triple-negative breast cancer cells inhibited cell proliferation and migration and promoted apoptosis (57). In a specific case of acute promyelocytic leukemia, formation of a hSSB2-RARA (retinoic acid receptor) chimera protein was linked to increased cancer predisposition (58). These results indicate that, given the non-essential nature of hSSB2 in the absence of stress, together with its prominent roles in genome repair, the targeting of its functions – among them, LLPS – can be relevant for cancer therapy *via* sensitizing cancer cells to increase the efficiency of existing drugs or to overcome drug resistance.

## MATERIALS AND METHODS

### General reaction conditions

Measurements were performed at 25°C in standard LLPS buffer containing 25 mM Tris-HCl (pH 7.4), 50 mM KCl, 10 mM MgCl_2_ and 1 mM DTT, unless otherwise indicated.

### Plasmids used for protein preparation

Plasmids pET29a-hNABP1 (encoding C-terminally His-tagged hSSB2), pET29a-hNABP2 (hSSB1), pET28a-C9ORF80 (INIP), and pGEX-6P-1-INTS3-FL (INTS3) were gifts from Yuliang Wu (Addgene plasmids #128307, 128306, 128418, and 128415, respectively) (10,12).

pET29a-hNABP2 encodes a hSSB1 protein variant harboring a Y85C amino acid substitution; by using QuikChange (Agilent) mutagenesis kit we changed its sequence to encode wild-type hSSB1.

A plasmid encoding hSSB2 fused to an N-terminal His-tag and TEV protease cleavage site was generated from pET29a-hNABP1 (hSSB2). A STOP codon was introduced after the triplet encoding the last amino acid of hSSB2. In addition, a region encoding the histidine tag and a TEV cleavage site was cloned in frame between the START codon and the first triplet encoding hSSB2.

The plasmid encoding the IDR-truncated hSSB2 variant was generated from the plasmid encoding N-terminally His-tagged hSSB2 by introducing a STOP codon after the triplet for amino acid residue 100 of hSSB2.

Plasmids encoding *E. coli* SSB, human BLM helicase, human Topoisomerase IIIa, human RMI1 and RMI2 and EGFP are described in (32,49).

### Protein expression and purification

C-terminally His-tagged hSSB2 was expressed and purified as described in (10) with the following modifications. Cell lysate obtained from hSSB2 expressing *E. coli* Rosetta 2 cells was loaded onto Ni-NTA beads equilibrated with buffer A (25 mM Tris-HCl pH 8.0, 500 mM NaCl, 10 % glycerol, 0.1 % TWEEN 20). The column was then washed with 20 column volumes of buffer B (25 mM Tris-HCl pH 8.0, 50 mM NaCl, 10 % glycerol, 0.1 % TWEEN 20, 50 mM imidazole). The protein was eluted with buffer B containing 250 mM imidazole. Eluted protein was concentrated using Amicon Ultra 10K spin columns (Sigma-Aldrich) and dialyzed against hSSB storage buffer (25 mM Tris-HCl pH 8.0, 50 mM KCl, 100 mM MgCl_2_, 10 % glycerol, 1 mM DTT).

hSSB2-dIDR and N-terminally His-tagged hSSB2 were purified as described above for the C-terminally His-tagged protein.

Histidine tag-free hSSB2 was prepared from N-terminally His-tagged hSSB2 using TEV protease at 1:500 TEV:hSSB2 molar ratio. Digestion was performed overnight at room temperature. The sample was passed through a Ni-NTA column equilibrated with 25 mM Tris-HCl pH 8.0, 50 mM NaCl, 50 mM imidazole, 0.1 % TWEEN 20, 10 % glycerol, and 20 mM MgCl_2_, and the flowthrough containing hSSB2 was collected.

hSSB1 was purified as described above for hSSB2, with the exception that the protein eluted from the Ni-NTA column was loaded on HiTrap Heparin column (GE Healthcare) equilibrated with HP1 buffer (25 mM Tris-HCl pH 8.0, 50 mM NaCl, 10 % glycerol). The column was washed with 2 column volumes of HP1 buffer to remove imidazole, and then eluted with HP2 buffer (25 mM Tris-HCl pH 8.0, 1 M NaCl, 10% glycerol). The eluted protein was concentrated using Amicon Ultra 10K spin columns (Sigma-Aldrich) and dialyzed against hSSB storage buffer.

INTS3 and INIP were purified as described in (12) with the following modifications for INTS3. After cell lysis by sonication, GST-tagged INTS3 was loaded onto a Glutathione Agarose (Pierce™) column equilibrated with buffer A (25 mM Tris-HCl pH 8.0, 500 mM NaCl, 1 mM EDTA, 1 mM DTT, 0.2 % TWEEN 20, 10 % glycerol). The column was washed with 2 column volumes of buffer A and then with 2 column volumes of buffer B (25 mM Tris-HCl pH 8.0, 150 mM NaCl, 1 mM DTT, 0.2 % TWEEN 20, 10 % glycerol). Precision protease (10 Unit/ml, Sigma-Aldrich) in buffer B was loaded onto the column, and INTS3 was digested overnight. Protein was eluted with buffer B, and fractions were pooled and concentrated using Amicon Ultra 100K spin columns (Sigma-Aldrich), The concentrated sample was dialyzed against storage buffer (25 mM Tris-HCl pH 8.0, 50 mM NaCl, 10 % glycerol, 1 mM DTT).

Human BLM, the TRR complex, *E. coli* SSB and EGFP were prepared as described in (32,49).

Purity of protein preparations was checked by SDS-PAGE. Protein concentrations were determined using the Bradford method. Purified proteins were frozen into droplets and stored in liquid N_2_.

### Fluorescent labeling of proteins

hSSB2 was labeled on intrinsic cysteines using 5-IAF dye (5-iodoacetamido-fluorescein, Thermo Fisher). Before labeling, DTT was removed from hSSB2 preparations using a PD10 (GE Healthcare) gel filtration column. IAF was introduced at 1.5-fold molar excess over protein. The labeling reaction was carried out for 3.5 h at room temperature under argon gas. Separation of the labeled protein from residual dye was carried out using a PD10 (GE Healthcare) gel filtration column pre-equilibrated with hSSB2 storage buffer, and residual dye was further removed by dialysis against storage buffer.

INIP and the TRR complex were labeled on N-termini using AF488 dye (Alexa Fluor 488 carboxylic acid succinimidyl ester, Thermo Fisher). Before labeling, the storage buffer was exchanged to labeling buffers (INIP: 50 mM MES pH 6.5, 50 mM KCl, 10 % glycerol; TRR: 50 mM HEPES pH 7.5, 200 mM NaCl, 1 mM DTT) by dialysis. AF488 was mixed in 4-fold molar excess over INIP and 15-fold molar excess over the TRR complex, and the solutions were incubated for 5 h at room temperature for INIP, and at 4°C for TRR. Reactions were stopped by the addition of Tris-HCl (50 mM final concentration). Labeled INIP was purified using a PD10 (GE Healthcare) gel filtration column pre-equilibrated with INIP storage buffer (25 mM Tris-HCl pH 8.0, 150 mM KCl, 10 % glycerol). The protein was dialyzed against storage buffer and repeatedly filtrated using Amicon Ultra 10K spin columns to remove residual free dye. Labeled TRR was dialyzed against its storage buffer (50 mM Tris-HCl pH 7.5, 200 mM NaCl, 10 % glycerol, 1 mM DTT).

hSSB1 and INTS3 proteins were labeled using AF647 dye (Alexa Fluor 647 carboxylic acid succinimidyl ester, Thermo Fisher). The storage buffer was exchanged by dialysis to labeling buffer containing 50 mM MES pH 6.5, 50 mM KCl, 100 mM MgCl_2_, 5 mM DTT. AF647 was applied at 1.5-fold molar excess over proteins. Labeling reactions were carried out for 3.5 h. Reactions were then stopped by the addition of Tris-HCl (to 50 mM final concentration). Labeled proteins were purified with a PD10 (GE Healthcare) gel filtration column pre-equilibrated with the storage buffers of proteins (hSSB1: 25 mM Tris-HCl pH 8.0, 50 mM NaCl, 100 mM MgCl_2_, 10 % glycerol, 1 mM DTT; INTS3: 25 mM Tris-HCl pH 8.0, 50 mM NaCl, 10 % glycerol, 1 mM DTT). Samples were dialyzed against their respective storage buffers and, in the case of INTS3, an extra ultrafiltration step (Amicon Ultra 100K spin column (Sigma-Aldrich)) was applied to fully remove residual dye.

Protein concentrations were determined by Bradford method. Dye concentrations were determined by visible light spectrometry using *ε*_494_ = 80,000 M^-1^cm^-1^ for 5-IAF, ε_488_ = 73,000 M^−1^cm^−1^ for AF488, and ε_647_ = 270,000 M^−1^cm^−1^ for AF647. Labeling ratios were 60 % for hSSB2, 17 % for INIP, 10 % for hSSB1, 9.6 % for INTS3, and 143 % for the TRR complex (47 % of the total N-termini within the complex). Purity of proteins was checked by SDS-PAGE. All constructs were frozen in liquid N_2_ in small aliquots and stored at −80°C. BLM and EcSSB were labeled as in (32).

### Fluorescence anisotropy titrations

Fluorescence polarization of 15-μl samples was measured in 384-well low-volume nontransparent microplates (Greiner Bio-one, PN:784900) at 25°C in a Synergy H4 Hybrid Multi-Mode Microplate Reader (BioTek) and converted to anisotropy values. Measurements were carried out in standard LLPS buffer omitting MgCl_2_. 10 nM of 3’-fluorescein-labeled 36- mer ssDNA (ATTTTTGCGGATGGCTTAGAGCTTAATTGCGCAACG-fluorescein); 36-mer ssRNA (AUUUUUGCGGAUGGCUUAGAGCUUAAUUGCGCAACG-fluorescein); or 54-mer ssDNA (TCCTTTTGATAAGAGGTCATTTTTGCGGATGGCTTAGAGCTTAATTGCGCAACG-Fluorescein) oligonucleotides were used. Data were analyzed by fits using the Hill equation. Data analysis and visualization were performed using OriginLab 8.0 (Microcal corp.).

### Electrophoretic mobility shift assays (EMSA)

50 nM 5’-Cy3-labeled, 41-nt ssRNA (5’-Cy3-U41) or 41-nt ssDNA (5’-Cy3-dT41) were titrated with increasing concentrations of hSSB2 (0–30 µM) in standard LLPS buffer. Samples were run on a 1 % w/v agarose gel at 120 V for 20 minutes. Labeled nucleic acids were detected upon excitation with a 532-nm laser in a Typhoon Trio+ Variable Mode Imager (Amersham Biosciences). Data were analyzed by fits using the Hill equation. Pixel densitometry was performed using GelQuant Pro software v12 (DNR Bio Imaging Ltd.).

### Turbidity measurements

Turbidity (light depletion) of samples (40 µl) was measured at 600 nm wavelength in transparent 384-well plates (Nunc Thermo Fisher PN:242757) using a Tecan Infinite Nano plate reader instrument at 25°C. Measurements were performed in standard LLPS buffer, and hSSB2 was used at 20 µM concentration.

### Determination of the minimal ssDNA length required for efficient condensation

Data shown in **Fig. 2H** (ssDNA length dependence of log-transformed half-maximal ssDNA concentrations required for hSSB2 condensation) were analyzed using **Equation 1** to estimate the minimal ssDNA length required for efficient hSSB2 condensation:

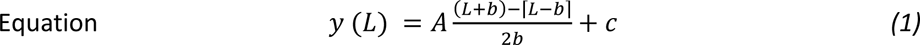

where *y* (*L*) is the logarithm of the half-maximal concentration determined using an ssDNA molecule comprising *L* nucleotides (determined from experiments in **Fig. 2G**), *A* is the amplitude, *b* is the length of ssDNA in nucleotides at which the function reaches saturation, and *c* is the y-axis intercept at *L* = 0.

We originally developed **Equation 1** to estimate the occluded site size of EcRecQ helicase based on the apparent dissociation constants obtained from ssDNA length dependent ATPase measurements (59). The model assumes that the change in standard Gibbs free energy (proportional to the logarithm of the dissociation constant) for protein.ssDNA interactions is linearly proportional to the length of ssDNA (assuming a uniform energetic contribution of each nucleotide that specifically interacts with the protein) as long as the length of ssDNA is smaller than the enzyme’s occluded site size, and remains constant when the ssDNA length exceeds the occluded site size (59). Since ssDNA binding triggers hSSB2 condensation, we can assume that the ssDNA concentration dependent turbidity signal observed in titration experiments (**Fig. 2G**) is proportional to the concentration of phase separation-prone hSSB2-ssDNA complexes. The amount of such complexes is determined by the law of mass action; thus, by the hSSB2 and ssDNA concentrations, and by the dissociation constant of the hSSB2-ssDNA interaction. The dissociation constant is assumed to depend on the length of ssDNA, at least partially due to similar reasons described above for EcRecQ. Accordingly, the logarithm of ssDNA concentrations required for half-maximal signal saturation showed a linear dependence on the ssDNA length until reaching a critical length, and remained constant afterwards.

### Epifluorescence microscopy

A Nikon Eclipse Ti-E TIRF microscope was used with apo TIRF 100x oil immersion objective (numerical aperture (NA) = 1.49) in epifluorescence mode. For fluorescence excitation, a 642-nm laser (56RCS/S2799, Melles Griot), a 543-nm laser (25-LGP-193–230, Melles Griot) and a Cyan 488-nm laser (Coherent) were used according to the excitation spectra of the applied fluorescent probes. Fluorescence was deflected to a ZT405/488/561/640rpc BS dichroic mirror and recorded with a Zyla sCMOS (ANDOR) camera.

Samples (20-μl) were visualized in μ-Slide Angiogenesis (Ibidi) microscope slides at 25°C. Images were recorded using NIS-Elements AR (Advanced Research) 4.50.00 software, using 2x2 pixel binning and 200-ms laser exposure time during acquisition.

### Image processing for epifluorescence microscopy

Fiji (ImageJ) software was used to process and analyze raw microscopic images. Image stacks and montages were generated from results of the analyzed experimental sets to adjust the brightness and contrast of images equally, using the automatic detection algorithm of Fiji. For representative purposes, images were background corrected using the built-in rolling ball background correction algorithm, unless otherwise stated.

Droplet size analysis (**Fig. 6F-H**) was carried out as in (32). To quantify the extent of condensation, mean gray values and total droplet areas of the middle image sections were calculated as follows. A stack of 3 background uncorrected images was generated from 3 different fields of view obtained at the given condition. Due to uneven illumination at the edges of raw images, a square region (600 x 600 pixels) centered to each image was selected for analysis (termed ‘middle ROI’). The mean gray value for each region was calculated as the sum of pixel intensities divided by the total number of pixels using the Stack Fitter plugin, and the average of mean gray values was calculated from the three analyzed images. Total droplet area was determined from the same regions after defining a signal threshold to distinguish between the background and the signal using the built-in thresholding algorithm. The built-in particle analyzer algorithm of Fiji (smallest detected size was set to 0.2 μm^2^, circularity 0.1–1) was used to determine outline and area of fluorescents spots. Determined areas were summarized for each image, and results were averaged for three analyzed regions.

Calculation of Pearson’s coefficients was performed on three background corrected total view field images for each specified condition using the Coloc 2 plugin of Fiji. Resulting Pearson’s coefficients were averaged.

## ACKNOWLEDGEMENTS

This work was supported by the “Momentum” Program of the Hungarian Academy of Sciences (LP2011-006/2011), ELTE KMOP-4.2.1/B-10-2011-0002, NKFIH K-123989, and NKFIH K-134595 grants to M.K. The project was supported by the NRDIO [VEKOP-2.3.3-15- 2016-00007] grant to ELTE. G.M.H. was supported by the Premium Postdoctoral Program of the Hungarian Academy of Sciences [PREMIUM-2017-17 to G.M.H.]. Z.J.K. and J.P. were supported by the New National Excellence Program of the Ministry for Innovation and Technology [Grants ÚNKP-21-3 to Z.J.K. and ÚNKP-19-2 to J.P.], and the Co-operative Doctoral Program of the Ministry of Innovation and Technology financed from the National Research, Development and Innovation Fund. This work was completed in the ELTE “SzintPlusz” Thematic Excellence Programme supported by the Hungarian Ministry for Innovation and Technology, and also in the framework of Project no. 2018-1.2.1-NKP-2018-00005 implemented with the support provided from the National Research, Development and Innovation Fund of Hungary, financed under the 2018-1.2.1-NKP funding scheme. This project has received funding from the HUN-REN Hungarian Research Network.

## DATA AVAILABILITY

Correspondence and material requests should be addressed to Mihaly Kovacs.

## AUTHOR CONTRIBUTIONS

G.M.H. and M.K. conceived the project. Z.J.K., G.M.H., J.P., G.S., and M.K. designed the experiments. Z.J.K., J.P., N.K., J.H., H.H.P., L.M., and L.K. performed the experiments. Z.J.K., G.M.H., J.P., and M.K. analyzed the data. Z.J.K, G.M.H., and M.K. wrote the manuscript. All authors contributed to editing and revision of the manuscript.

## REFERENCES

1. Richard, D.J., Bolderson, E., Cubeddu, L., Wadsworth, R.I., Savage, K., Sharma, G.G., Nicolette, M.L., Tsvetanov, S., McIlwraith, M.J., Pandita, R.K. et al. (2008) Single-stranded DNA-binding protein hSSB1 is critical for genomic stability. Nature, 453, 677–681.

2. Kang, H.S., Beak, J.Y., Kim, Y.S., Petrovich, R.M., Collins, J.B., Grissom, S.F. and Jetten, A.M. (2006) NABP1, a novel RORgamma-regulated gene encoding a single-stranded nucleic-acid-binding protein. Biochem J, 397, 89–99.

3. Ashton, N.W., Bolderson, E., Cubeddu, L., O’Byrne, K.J. and Richard, D.J. (2013) Human single-stranded DNA binding proteins are essential for maintaining genomic stability. BMC Mol Biol, 14, 9.

4. Wu, Y., Lu, J. and Kang, T. (2016) Human single-stranded DNA binding proteins: guardians of genome stability. Acta Biochim Biophys Sin (Shanghai), 48, 671–677.

5. Feldhahn, N., Ferretti, E., Robbiani, D.F., Callen, E., Deroubaix, S., Selleri, L., Nussenzweig, A. and Nussenzweig, M.C. (2012) The hSSB1 orthologue Obfc2b is essential for skeletogenesis but dispensable for the DNA damage response in vivo. EMBO J, 31, 4045–4056.

6. Shi, W., Vu, T., Boucher, D., Biernacka, A., Nde, J., Pandita, R.K., Straube, J., Boyle, G.M., Al-Ejeh, F., Nag, P. et al. (2017) Ssb1 and Ssb2 cooperate to regulate mouse hematopoietic stem and progenitor cells by resolving replicative stress. Blood, 129, 2479–2492.

7. Shi, W., Bain, A.L., Schwer, B., Al-Ejeh, F., Smith, C., Wong, L., Chai, H., Miranda, M.S., Ho, U., Kawaguchi, M. et al. (2013) Essential developmental, genomic stability, and tumour suppressor functions of the mouse orthologue of hSSB1/NABP2. PLoS Genet, 9, e1003298.

8. Croft, L.V., Bolderson, E., Adams, M.N., El-Kamand, S., Kariawasam, R., Cubeddu, L., Gamsjaeger, R. and Richard, D.J. (2019) Human single-stranded DNA binding protein 1 (hSSB1, OBFC2B), a critical component of the DNA damage response. Semin Cell Dev Biol, 86, 121–128.

9. Pfeifer, M., Brem, R., Lippert, T.P., Boulianne, B., Ho, H.N., Robinson, M.E., Stebbing, J. and Feldhahn, N. (2019) SSB1/SSB2 Proteins Safeguard B Cell Development by Protecting the Genomes of B Cell Precursors. J Immunol, 202, 3423–3433.

10. Vidhyasagar, V., He, Y., Guo, M., Ding, H., Talwar, T., Nguyen, V., Nwosu, J., Katselis, G. and Wu, Y. (2016) C-termini are essential and distinct for nucleic acid binding of human NABP1 and NABP2. Biochim Biophys Acta, 1860, 371–383.

11. Skaar, J.R., Ferris, A.L., Wu, X., Saraf, A., Khanna, K.K., Florens, L., Washburn, M.P., Hughes, S.H. and Pagano, M. (2015) The Integrator complex controls the termination of transcription at diverse classes of gene targets. Cell Res, 25, 288–305.

12. Vidhyasagar, V., He, Y., Guo, M., Talwar, T., Singh, R.S., Yadav, M., Katselis, G., Vizeacoumar, F.J., Lukong, K.E. and Wu, Y. (2018) Biochemical characterization of INTS3 and C9ORF80, two subunits of hNABP1/2 heterotrimeric complex in nucleic acid binding. Biochem J, 475, 45–60.

13. Li, Y., Bolderson, E., Kumar, R., Muniandy, P.A., Xue, Y., Richard, D.J., Seidman, M., Pandita, T.K., Khanna, K.K. and Wang, W. (2009) HSSB1 and hSSB2 form similar multiprotein complexes that participate in DNA damage response. J Biol Chem, 284, 23525–23531.

14. Boucher, D., Kariawasam, R., Burgess, J., Gimenez, A., Ocampo, T.E., Ferguson, B., Naqi, A., Walker, G.J., Bolderson, E., Gamsjaeger, R. et al. (2021) hSSB2 (NABP1) is required for the recruitment of RPA during the cellular response to DNA UV damage. Sci Rep, 11, 20256.

15. Boucher, D., Vu, T., Bain, A.L., Tagliaro-Jahns, M., Shi, W., Lane, S.W. and Khanna, K.K. (2015) Ssb2/Nabp1 is dispensable for thymic maturation, male fertility, and DNA repair in mice. FASEB J, 29, 3326–3334.

16. Gu, P., Deng, W., Lei, M. and Chang, S. (2013) Single strand DNA binding proteins 1 and 2 protect newly replicated telomeres. Cell Res, 23, 705–719.

17. Vernin, C., Thenoz, M., Pinatel, C., Gessain, A., Gout, O., Delfau-Larue, M.H., Nazaret, N., Legras-Lachuer, C., Wattel, E. and Mortreux, F. (2014) HTLV-1 bZIP factor HBZ promotes cell proliferation and genetic instability by activating OncomiRs. Cancer Res, 74, 6082–6093.

18. Bain, A.L., Shi, W. and Khanna, K.K. (2013) Mouse models uncap novel roles of SSBs. Cell Res, 23, 744–745.

19. Ashton, N.W., Loo, D., Paquet, N., O’Byrne, K.J. and Richard, D.J. (2016) Novel insight into the composition of human single-stranded DNA-binding protein 1 (hSSB1)-containing protein complexes. BMC Mol Biol, 17, 24.

20. Croft, L.V., Ashton, N.W., Paquet, N., Bolderson, E., O’Byrne, K.J. and Richard, D.J. (2017) hSSB1 associates with and promotes stability of the BLM helicase. BMC Mol Biol, 18, 13.

21. Xu, S., Feng, Z., Zhang, M., Wu, Y., Sang, Y., Xu, H., Lv, X., Hu, K., Cao, J., Zhang, R. et al. (2011) hSSB1 binds and protects p21 from ubiquitin-mediated degradation and positively correlates with p21 in human hepatocellular carcinomas. Oncogene, 30, 2219–2229.

22. Richard, D.J., Cubeddu, L., Urquhart, A.J., Bain, A., Bolderson, E., Menon, D., White, M.F. and Khanna, K.K. (2011) hSSB1 interacts directly with the MRN complex stimulating its recruitment to DNA double-strand breaks and its endo-nuclease activity. Nucleic Acids Res, 39, 3643–3651.

23. Zhang, F., Wu, J. and Yu, X. (2009) Integrator3, a partner of single-stranded DNA-binding protein 1, participates in the DNA damage response. J Biol Chem, 284, 30408–30415.

24. Huang, J., Gong, Z., Ghosal, G. and Chen, J. (2009) SOSS complexes participate in the maintenance of genomic stability. Mol Cell, 35, 384–393.

25. Skaar, J.R., Richard, D.J., Saraf, A., Toschi, A., Bolderson, E., Florens, L., Washburn, M.P., Khanna, K.K. and Pagano, M. (2009) INTS3 controls the hSSB1-mediated DNA damage response. J Cell Biol, 187, 25–32.

26. Richard, D.J., Bolderson, E. and Khanna, K.K. (2009) Multiple human single-stranded DNA binding proteins function in genome maintenance: structural, biochemical and functional analysis. Crit Rev Biochem Mol Biol, 44, 98–116.

27. Lawson, T., El-Kamand, S., Boucher, D., Duong, D.C., Kariawasam, R., Bonvin, A., Richard, D.J., Gamsjaeger, R. and Cubeddu, L. (2020) The structural details of the interaction of single-stranded DNA binding protein hSSB2 (NABP1/OBFC2A) with UV-damaged DNA. Proteins, 88, 319–326.

28. Paquet, N., Adams, M.N., Ashton, N.W., Touma, C., Gamsjaeger, R., Cubeddu, L., Leong, V., Beard, S., Bolderson, E., Botting, C.H. et al. (2016) hSSB1 (NABP2/OBFC2B) is regulated by oxidative stress. Sci Rep, 6, 27446.

29. Paquet, N., Adams, M.N., Leong, V., Ashton, N.W., Touma, C., Gamsjaeger, R., Cubeddu, L., Beard, S., Burgess, J.T., Bolderson, E. et al. (2015) hSSB1 (NABP2/ OBFC2B) is required for the repair of 8-oxo-guanine by the hOGG1-mediated base excision repair pathway. Nucleic Acids Res, 43, 8817–8829.

30. Rajapakse, A., Suraweera, A., Boucher, D., Naqi, A., O’Byrne, K., Richard, D.J. and Croft, L.V. (2020) Redox Regulation in the Base Excision Repair Pathway: Old and New Players as Cancer Therapeutic Targets. Curr Med Chem, 27, 1901–1921.

31. Lawson, T., El-Kamand, S., Kariawasam, R., Richard, D.J., Cubeddu, L. and Gamsjaeger, R. (2019) A Structural Perspective on the Regulation of Human Single-Stranded DNA Binding Protein 1 (hSSB1, OBFC2B) Function in DNA Repair. Comput Struct Biotechnol J, 17, 441–446.

32. Harami, G.M., Kovacs, Z.J., Pancsa, R., Palinkas, J., Barath, V., Tarnok, K., Malnasi-Csizmadia, A. and Kovacs, M. (2020) Phase separation by ssDNA binding protein controlled via protein-protein and protein-DNA interactions. Proc Natl Acad Sci U S A, 117, 26206–26217.

33. Hyman, A.A., Weber, C.A. and Julicher, F. (2014) Liquid-liquid phase separation in biology. Annu Rev Cell Dev Biol, 30, 39–58.

34. Gao, Z., Zhang, W., Chang, R., Zhang, S., Yang, G. and Zhao, G. (2021) Liquid-Liquid Phase Separation: Unraveling the Enigma of Biomolecular Condensates in Microbial Cells. Front Microbiol, 12, 751880.

35. Li, H., Ernst, C., Kolonko-Adamska, M., Greb-Markiewicz, B., Man, J., Parissi, V. and Ng, B.W. (2022) Phase separation in viral infections. Trends Microbiol, 30, 1217–1231.

36. Wang, B., Zhang, L., Dai, T., Qin, Z., Lu, H., Zhang, L. and Zhou, F. (2021) Liquid-liquid phase separation in human health and diseases. Signal Transduct Target Ther, 6, 290.

37. Zhao, T., Liu, Y., Wang, Z., He, R., Xiang Zhang, J., Xu, F., Lei, M., Deci, M.B., Nguyen, J. and Bianco, P.R. (2019) Super-resolution imaging reveals changes in Escherichia coli SSB localization in response to DNA damage. Genes Cells, 24, 814–826.

38. Spegg, V., Panagopoulos, A., Stout, M., Krishnan, A., Reginato, G., Imhof, R., Roschitzki, B., Cejka, P. and Altmeyer, M. (2023) Phase separation properties of RPA combine high-affinity ssDNA binding with dynamic condensate functions at telomeres. Nat Struct Mol Biol, 30, 451–462.

39. Harami, G.M., Palinkas, J., Kovacs, Z.J., Tarnok, K., Harami-Papp, H., Hegedus, J., Mahmudova, L., Kucsma, N., Toth, S., Szakacs, G., et al. (2023) Redox-dependent condensation and cytoplasmic granulation by human ssDNA binding protein 1 delineate roles in oxidative stress response. bioRxiv, doi: 10.1101/2023.1107.1125.550517, PREPRINT: NOT PEER REVIEWED.

40. Hellman, L.M. and Fried, M.G. (2007) Electrophoretic mobility shift assay (EMSA) for detecting protein-nucleic acid interactions. Nat Protoc, 2, 1849–1861.

41. Alberti, S., Gladfelter, A. and Mittag, T. (2019) Considerations and Challenges in Studying Liquid-Liquid Phase Separation and Biomolecular Condensates. Cell, 176, 419–434.

42. Krainer, G., Welsh, T.J., Joseph, J.A., Espinosa, J.R., Wittmann, S., de Csillery, E., Sridhar, A., Toprakcioglu, Z., Gudiskyte, G., Czekalska, M.A., et al. (2021) Reentrant liquid condensate phase of proteins is stabilized by hydrophobic and non-ionic interactions. Nat Commun, 12, 1085.

43. Bootman, M.D. and Bultynck, G. (2020) Fundamentals of Cellular Calcium Signaling: A Primer. Cold Spring Harb Perspect Biol, 12.

44. Milo, R., Jorgensen, P., Moran, U., Weber, G. and Springer, M. (2010) BioNumbers--the database of key numbers in molecular and cell biology. Nucleic Acids Res, 38, D750–753.

45. Mangia, S., Giove, F. and Dinuzzo, M. (2012) Metabolic pathways and activity-dependent modulation of glutamate concentration in the human brain. Neurochem Res, 37, 2554–2561.

46. Bizard, A.H. and Hickson, I.D. (2020) The many lives of type IA topoisomerases. J Biol Chem, 295, 7138–7153.

47. Bizard, A.H. and Hickson, I.D. (2014) The dissolution of double Holliday junctions. Cold Spring Harb Perspect Biol, 6, a016477.

48. Bythell-Douglas, R. and Deans, A.J. (2021) A Structural Guide to the Bloom Syndrome Complex. Structure, 29, 99–113.

49. Harami, G.M., Palinkas, J., Seol, Y., Kovacs, Z.J., Gyimesi, M., Harami-Papp, H., Neuman, K.C. and Kovacs, M. (2022) The toposiomerase IIIalpha-RMI1-RMI2 complex orients human Bloom’s syndrome helicase for efficient disruption of D-loops. Nat Commun, 13, 654.

50. Ren, W., Chen, H., Sun, Q., Tang, X., Lim, S.C., Huang, J. and Song, H. (2014) Structural basis of SOSS1 complex assembly and recognition of ssDNA. Cell Rep, 6, 982–991.

51. Kozlov, A.G., Cheng, X., Zhang, H., Shinn, M.K., Weiland, E., Nguyen, B., Shkel, I.A., Zytkiewicz, E., Finkelstein, I.J., Record, M.T., Jr., et al. (2022) How Glutamate Promotes Liquid-liquid Phase Separation and DNA Binding Cooperativity of E. coli SSB Protein. J Mol Biol, 434, 167562.

52. Bekker-Jensen, D.B., Kelstrup, C.D., Batth, T.S., Larsen, S.C., Haldrup, C., Bramsen, J.B., Sorensen, K.D., Hoyer, S., Orntoft, T.F., Andersen, C.L. et al. (2017) An Optimized Shotgun Strategy for the Rapid Generation of Comprehensive Human Proteomes. Cell Syst, 4, 587–599 e584.

53. Maul, G.G. and Deaven, L. (1977) Quantitative determination of nuclear pore complexes in cycling cells with differing DNA content. J Cell Biol, 73, 748–760.

54. Andre, A.A.M. and Spruijt, E. (2020) Liquid-Liquid Phase Separation in Crowded Environments. Int J Mol Sci, 21.

55. Par, S., Vaides, S., VanderVere-Carozza, P.S., Pawelczak, K.S., Stewart, J. and Turchi, J.J. (2021) OB-Folds and Genome Maintenance: Targeting Protein-DNA Interactions for Cancer Therapy. Cancers (Basel), 13.

56. Adams, M.N., Croft, L.V., Urquhart, A., Saleem, M.A.M., Rockstroh, A., Duijf, P.H.G., Thomas, P.B., Ferguson, G.P., Najib, I.M., Shah, E.T. et al. (2023) hSSB1 (NABP2/OBFC2B) modulates the DNA damage and androgen-induced transcriptional response in prostate cancer. Prostate, 83, 628–640.

57. Wu, Q., Tang, X., Zhu, W., Li, Q., Zhang, X. and Li, H. (2021) The Potential Prognostic Role of Oligosaccharide-Binding Fold-Containing Protein 2A (OBFC2A) in Triple-Negative Breast Cancer. Front Oncol, 11, 751430.

58. Won, D., Shin, S.Y., Park, C.J., Jang, S., Chi, H.S., Lee, K.H., Lee, J.O. and Seo, E.J. (2013) OBFC2A/RARA: a novel fusion gene in variant acute promyelocytic leukemia. Blood, 121, 1432–1435.

59. Harami, G.M., Seol, Y., In, J., Ferencziova, V., Martina, M., Gyimesi, M., Sarlos, K., Kovacs, Z.J., Nagy, N.T., Sun, Y. et al. (2017) Shuttling along DNA and directed processing of D-loops by RecQ helicase support quality control of homologous recombination. Proc Natl Acad Sci U S A, 114, E466–E475.

