## Supplementary Material for "DNA-dependent phase separation by human SSB2 (NABP1/OBFC2A) protein points to adaptations to eukaryotic genome repair processes"


#### SUPPLEMENTARY FIGURES

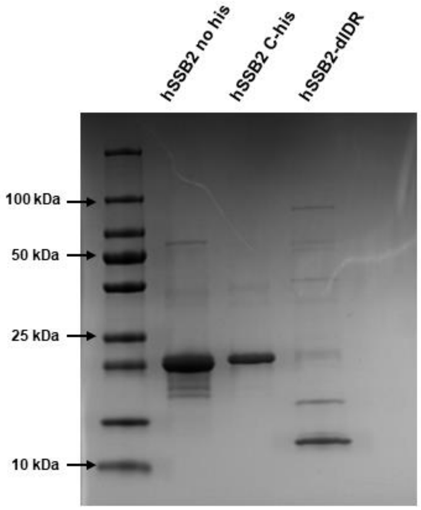


**Supplementary Figure 1. SDS-PAGE electrophoretograms of hSSB2 protein constructs.**
Coomassie-stained SDS-PAGE electrophoretograms of purified hSSB2, His-tagged hSSB2 and His-tagged hSSB2-dIDR protein constructs are shown. 5 µg protein was loaded for each construct. Calculated molecular weights of proteins were 23.4, 22.7 and 11.9 kDa, respectively. 4–20 % precast SDS-polyacrylamide gel (BIO-RAD #4561095) and PageRuler Plus Prestained Protein Ladder were used in the experiment.

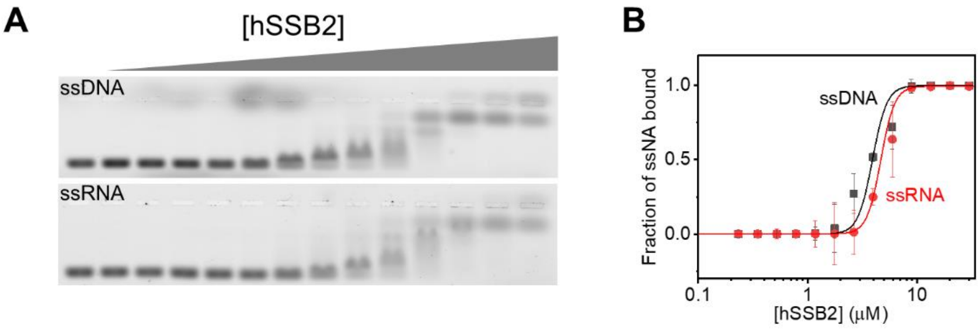


**Supplementary Figure 2. Nucleic acid binding to hSSB2 measured in EMSA assay.**
**(A)** Example electrophoretograms of electrophoretic mobility shift assays (EMSA). Constant concentration of 50 nM ssDNA (Cy3-labeled 41mer homopolymeric deoxythymidine; Cy3-labeled dT41) or ssRNA (Cy3-labeled 41mer homopolymeric uridine, Cy3-labeled U41) oligonucleotides were titrated with 0–30 µM hSSB2 in standard LLPS buffer. Samples were separated on a 1 w/v% agarose gel and visualized using a fluorescent gel scanner. The mobility of nucleic acids decreased upon hSSB2 binding. **(B)** Fractions of labeled ssDNA (black) and ssRNA (red) bound by hSSB2, analyzed by gel densitometry. Solid lines show best fits using the Hill equation. Means ± SD are shown for *n* = 2.

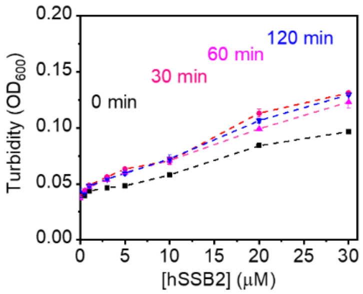


**Supplementary Figure 3. hSSB2 solutions contain a minor fraction of aggregated species in the absence of ssDNA.**
The turbidity of hSSB2 solutions is shown at indicated protein concentrations and incubation times in the absence on nucleic acids in standard LLPS buffer. hSSB2 stock solution was diluted into LLPS buffer and measurements were carried out immediately after mixing or after the indicated incubation times. Means ± SEM for *n* = 3 are shown. The concentration and time dependent increase in the turbidity signal indicates the presence and formation of particles that scatter light. The turbidity signal remained largely constant after 30 min. Accordingly, amorphous aggregates were detected in DIC microscopic measurements (**Fig. 2A**), indicating that hSSB2 is prone to aggregation at high concentrations under the applied conditions and in the absence of nucleic acids. Importantly, in the presence of ssDNA, hSSB2 forms LLPS droplets of liquid nature (**Fig. 2A-B**).

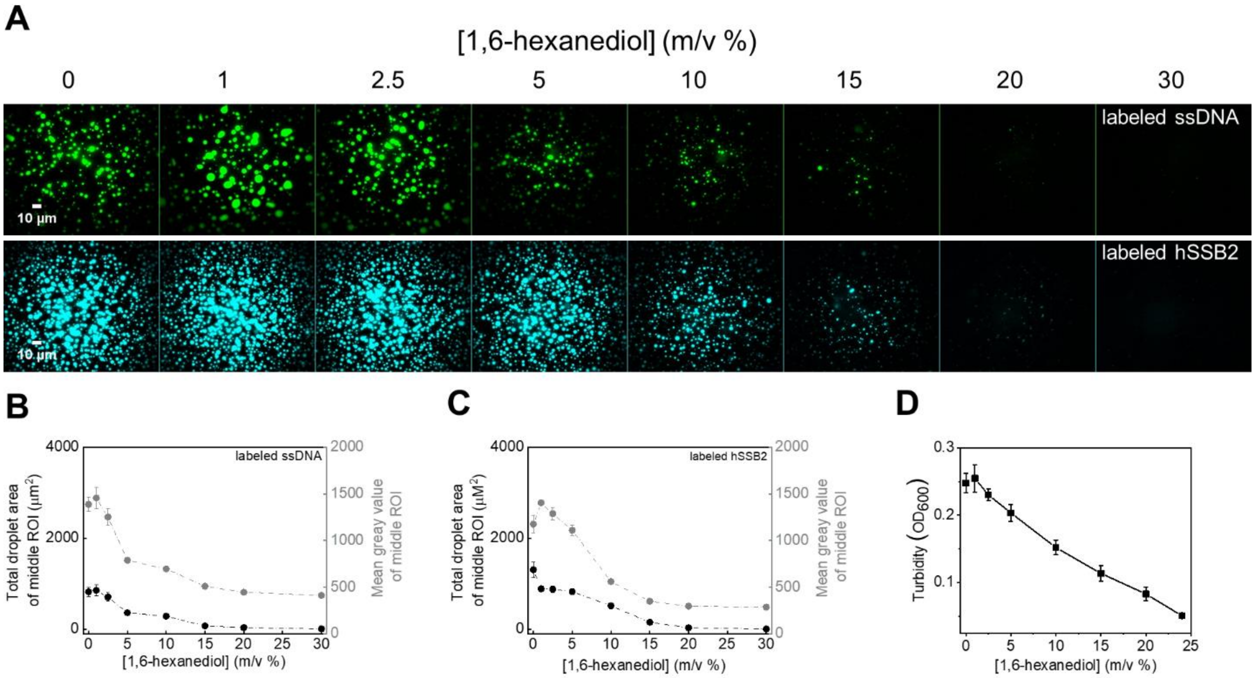


**Supplementary Figure 4. Inhibition of hSSB2 LLPS by 1,6-hexanediol.**

**(A)** Fluorescence microscopic images showing the inhibitory effect of the aliphatic alcohol 1,6-hexanediol on hSSB2 droplet formation. Upper row: 5 µM hSSB2, 2 µM dT79 containing 0.1 µM Cy3-labeled dT79. Bottom row: 10 µM hSSB2 containing 0.1 µM fluorescein-labeled hSSB2 and 1 µM dT79. Before imaging, samples were incubated in standard LLPS buffer containing indicated concentration of 1,6-hexanediol for 30 and 60 minutes (upper and lower rows), respectively.

**(B-C)** 1,6-hexanediol concentration dependence of the total droplet area and mean gray value of middle ROIs (see Methods) determined from panel **A**. Means ± SEM are shown for *n* = 3.

**(D)** Turbidity measurements performed upon mixing 15 µM unlabeled hSSB2 and 2 µM dT79 in standard LLPS buffer, containing indicated concentrations of 1,6-hexanediol. Means ± SD for *n* = 2 are shown.

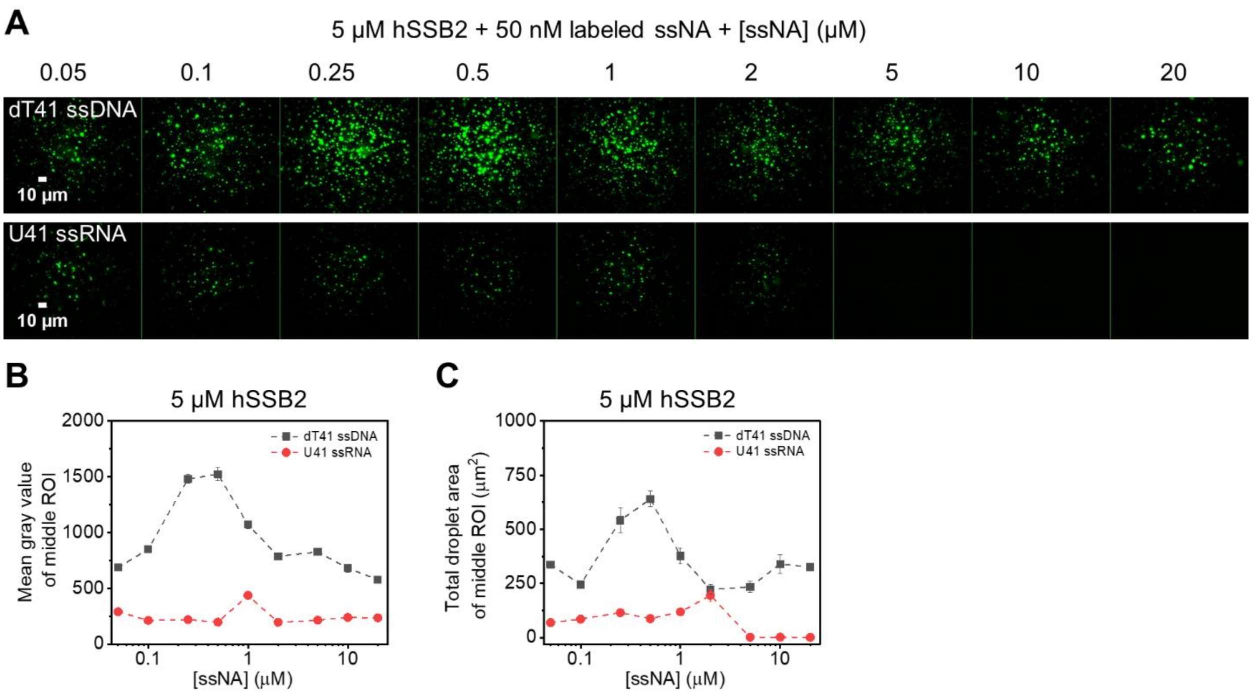


**Supplementary Figure 5. Differential effects of ssDNA and ssRNA on hSSB2 LLPS are also observed at subphysiological ion concentrations.**

**(A)** Fluorescence microscopic images recorded upon mixing 5 µM hSSB2, 50 nM Cy3-labeled dT41 ssDNA or U41 ssRNA, and indicated concentrations of unlabeled dT41 or U41, respectively. Samples were incubated in LLPS buffer omitting salts for 30 minutes before imaging.

**(B-C)** Mean gray value and total droplet area of middle ROIs determined from panel **A**. Means ± SEM are shown for *n* = 3.

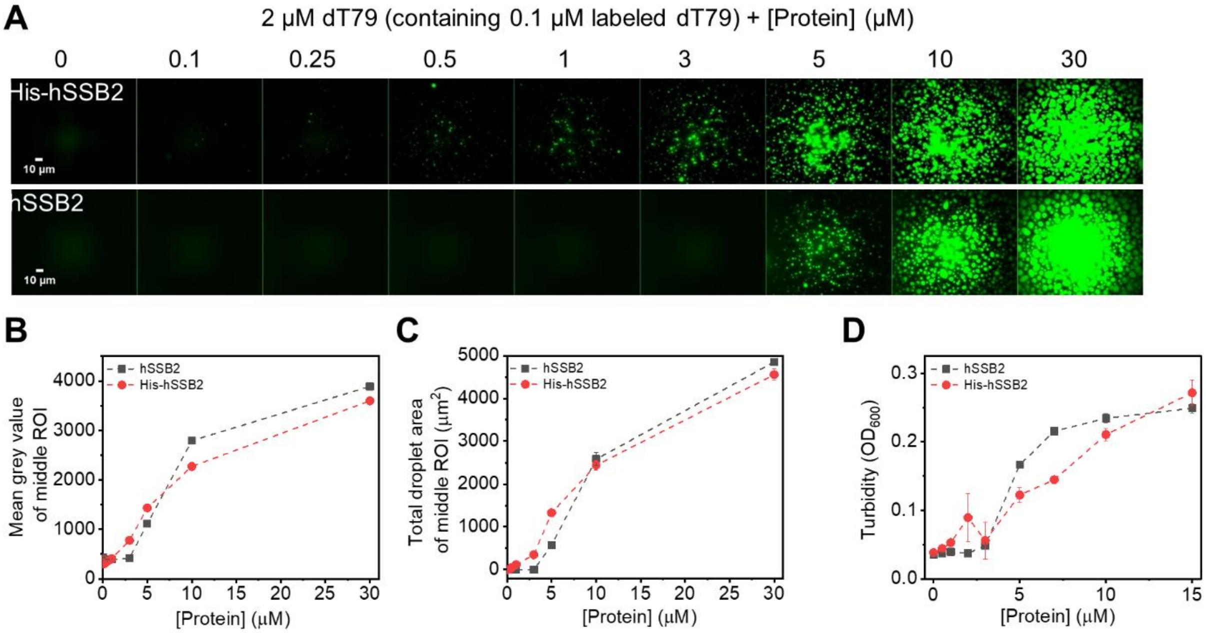


**Supplementary Figure 6. The presence of C-terminal histidine tag on hSSB2 moderately increase the LLPS propensity.**

**(A)** Fluorescence microscopic images of samples containing 2 μM dT79 (containing 0.1 μM Cy3-dT79) and indicated concentrations of proteins in standard LLPS buffer.

**(B-C)** Protein (tag-free hSSB2 (black), histidine-tagged hSSB2 (red)) concentration dependence of the total droplet area and mean gray value of middle ROIs determined (see Methods) from experiments in panel **A**. Means ± SEM are shown for *n* = 3.

**(D)** Turbidity measurements performed upon mixing increasing concentrations of unlabeled tag-free hSSB2 (black), or histidine-tagged hSSB2 (red) with 1 μM dT79 in standard LLPS buffer. Means ± SD for *n* = 2 are shown.

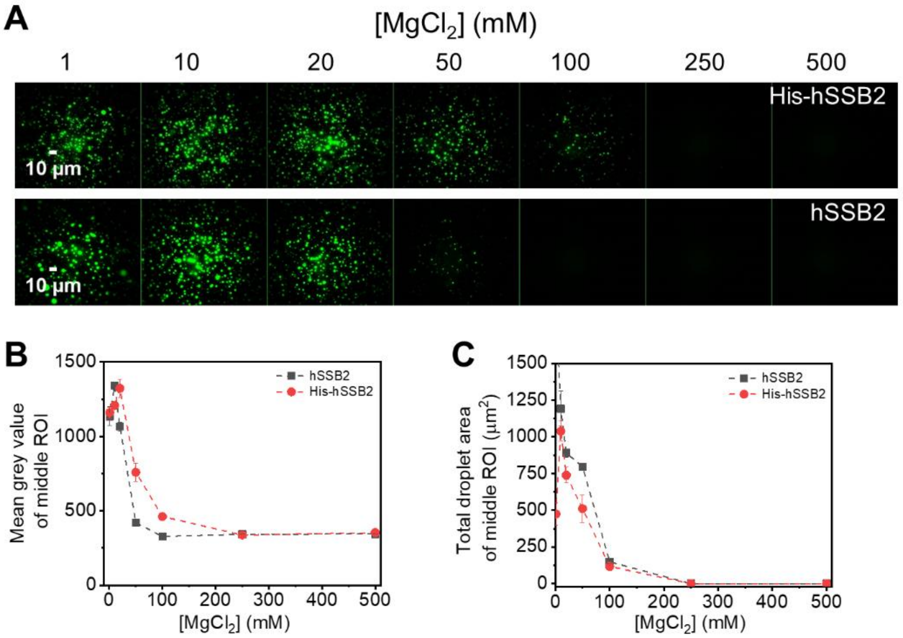


**Supplementary Figure 7. The presence of a C-terminal histidine tag on hSSB2 does not influence the sensitivity of hSSB2 LLPS to ionic conditions.**

**(A)** Representative fluorescence microscopic images of samples containing 5 µM histidine-tagged hSSB2 or tag-free hSSB2, 2 µM dT79 (containing 0.1 µM Cy3-dT79) in LLPS buffer containing 50 mM KCl and indicated concentration of $MgCl_2$. Samples were incubated for 30 minutes before imaging.

**(B-C)** Magnesium concentration dependence of the **(B)** mean gray value and **(C)** total droplet area of middle ROIs (see Methods) determined for histidine-tagged hSSB2 (red) and tag-free hSSB2 (black) in experiments shown in panel **A**. Means ± SEM are shown for *n* = 3.

### SUPPLEMENTARY TABLES

| Experiment | Nucleic acid | $K_d$ (µM) | *n* |
|---|---|---|---|
| Fluorescence anisotropy | flu-ss36 DNA | 0.53 ± 0.02 | 2.7 ± 0.3 |
| Fluorescence anisotropy | flu-ss36 RNA | 0.87 ± 0.17 | 3.3 ± 1.0 |
| EMSA | Cy3-dT41ssDNA | 3.9 ± 0.1 | 6.6 ± 1.7 |
| EMSA | Cy3-U41 ssRNA | 4.7 ± 0.6 | 6.9 ± 0.2 |

**Supplementary Table 1. ssDNA and ssRNA binding parameters of hSSB2 determined from fluorescence anisotropy titrations and electrophoretic mobility shift assays.**

Equilibrium dissociation constants ($K_d$) and Hill-coefficients (*n*) ± fitting errors are reported. Values were determined in by fitting the Hill equation to binding curves shown in **Fig. 1C** and **Supplementary Fig. 2**. Obtained Hill coefficient values are in line with hSSB2:ssDNA binding stoichiometries inferred from fluorescence microscopic LLPS assay results (**Fig. 2C-D**).

| Protein | Nucleic acid | $K_d$ (μM) | *n* |
|---|---|---|---|
| hSSB2 | ss54 ssDNA | 0.087 ± 0.007 | 1.1 ± 0.1 |
| hSSB2-dIDR | ss54 ssDNA | 0.85 ± 0.02 | 2.4 ± 0.2 |

**Supplementary Table 2. ssDNA binding parameters of hSSB2 and hSSB2-dIDR determined in fluorescence anisotropy titrations.** Equilibrium dissociation constants ($K_d$) and Hill-coefficients (*n*) ± fitting errors reported. Values were determined by fitting the Hill equation to binding curves shown in **Fig. 4B**.
